# The anticonvulsant and mood-stabilizing drug valproic acid attracts *C. elegans* and activates chemosensory neurons via a cGMP signaling pathway

**DOI:** 10.1101/2025.11.11.687732

**Authors:** Lucero E. Rogel-Hernandez, Helena Casademunt, Aravinthan D.T. Samuel, Miriam B. Goodman

**Affiliations:** Department of Molecular and Cellular Physiology, Stanford University, Stanford, CA 94304; Department of Physics and Center for Brain Science, Harvard University, Cambridge, MA 02134

## Abstract

Valproic acid (VPA) is a drug with both anticonvulsant and antimanic properties. It has been widely prescribed to treat epilepsy, bipolar disorder, and other neuropsychiatric conditions for decades, but prenatal exposure is linked to developmental anomalies, cognitive deficits, and an increased risk of autism. Remarkably, the molecular basis of VPA action in the brain is not fully understood. To determine VPA’s effect on the nervous system without requiring systemic drug application that induce toxic effects, we exploited the well-characterized chemotaxis behavior of the nematode *Caenorhabditis elegans*. We found that *C. elegans* are attracted to VPA, and this behavior is missing in animals lacking the *tax-4* ion channel and the *tax-4*-expressing AWC chemosensory neurons. To test the idea that VPA directly activates the AWC neurons, we performed calcium imaging studies in a line expressing GCaMP6s in all amphid chemosensory neurons in the head. We found that VPA evoked calcium transients consistently in the AWC neurons and variably in AWB and ASH. As cyclic nucleotide-gated channels, the active state of *tax-4* increases upon the binding of cGMP. Within the worm’s chemosensory nervous system, receptor guanylate cyclases (rGCs) facilitate the synthesis of cGMP and serve as chemoreceptors for various chemical cues. By performing chemotaxis assays and calcium imaging experiments with rGC mutants against VPA, we found that *odr-1* and *gcy-28* are essential for mediating VPA attraction, as their absence disrupts both behavior and control AWC calcium transients. However, given their broad use in chemosensation, *odr-1* and *gcy-28* most likely act as downstream effectors rather than as direct targets. These findings compelled us to investigate whether chemoreceptors belonging to the G protein-coupled receptor (GPCR) family contribute to VPA sensing. We approached this hypothesis by conducting chemotaxis assays with Gα mutants and identified several (*odr-3*, *egl-30*, *gpa-2;gpa-3*) whose absence leads to a loss of VPA attraction. Thus, future studies should focus on elucidating the mechanisms by which GPCRs contribute to VPA sensing and cGMP signaling.

## Introduction

Valproic acid (VPA) is a widely prescribed broad-spectrum antiepileptic drug and a mood stabilizer for bipolar disorder [1]. Although its mode of action in the nervous system remains elusive and unclear, it has been linked to increased GABA levels in the brain, an effect proposed to account for VPA’s antiepileptic properties [2]. Other candidate signaling mechanisms have been reported, including modulation of arachidonic acid and inositol levels in the brain and indirect inhibition of protein kinase C (PKC) and mitogen-activated protein kinase (MAPK) (Reviewed in [3]). Along with these established and emerging therapeutic benefits, VPA is linked to undesirable effects such as weight gain, hair loss, tremors, pancreatitis, and liver dysfunction [4]. Adverse effects arising from prenatal exposure are also significant: in humans and other mammals, studies have shown that prenatal exposure to this drug leads to developmental anomalies in offspring, such as neural tube, limb, and cardiac defects, and increases the risk of autism spectrum disorders [5]. Transgenerational effects might not be limited to mothers, since children born to VPA-treated fathers may also have an increased risk of neurodevelopmental conditions [6]. From this brief overview of a large literature, it is evident that improved clarity regarding the molecular targets of VPA in the nervous system might enable the development of safe and efficacious therapeutics to manage the neurological diseases treated with VPA today with an improved risk-reward profile.

The effect of VPA on neural function and behavior has been investigated in several invertebrates and *ex vivo* mammalian preparations [7–16], including *Caenorhabditis elegans* nematodes [17–20]. Wild-type *C. elegans* nematodes grown in the presence of VPA display defects in defecation, egg-laying, reduced brood size [17], as well as abnormal DNA methylation [20] and lifespan [18,19]. Whether in mammals or invertebrates, the toxic effects of whole-body exposure to VPA complicate investigations of how this drug affects the nervous system is uncertain. We reasoned that chemotaxis behavior could provide insight into this question without requiring whole-body exposure to the drug. To achieve this goal, we used *C. elegans* as a chemical detector and adapted our high-throughput chemotaxis-based screening platform [21] to evaluate VPA-induced chemotaxis. *C. elegans* relies on ciliated chemosensory neurons to perform chemotaxis. Each cilium is densely packed with a wide assortment of chemoreceptors that survey the extracellular space for chemical signals [22]. When a ligand activates a chemoreceptor, these proteins convert binding on their surfaces into electrical signals by relaying the information to downstream cyclic nucleotide-gated (CNG) or transient receptor potential (TRP) ion channels via the use of second messengers, such as cyclic nucleotides (cAMP, cGMP) or various lipid-derived mediators (i.e., DAG, IP3), respectively [22]. CNG or TRP channel activation induces membrane depolarization and initiates behavioral responses to attractant and repellant chemicals in the environment [22].

Chemoreception relies on a variety of receptor types, including receptor guanylate cyclases (rGCs) and G protein-coupled receptors (GPCRs), to enable the detection of environmental cues [23]. Combined, there are over 1,700 genes encoding these potential chemoreceptors in *C. elegans* [24–26]. This immense number and diversity of chemoreceptors [26] exceeds that found in vertebrate genomes [27] and is likely to enable *C. elegans* to detect a large number of environmental chemicals. In principle, each chemoreceptor can respond to multiple chemicals, and each chemical can activate multiple chemoreceptors, thus granting *C. elegans* the potential to detect and discriminate among thousands of chemical cues. One can speculate that having a large and diverse repertoire of chemoreceptors could allow for the detection of a more diverse panel of chemicals. Thus, *C. elegans* is well suited system to test a variety of ligands, including drugs like VPA. If such ligands are capable of being perceived by *C. elegans,* this creates an opportunity to explore ligand-receptor interactions and to characterize new binding sites, which in turn offers valuable insights into comparative biology that could help identify key human targets to advance biomedical research and drug development.

Combining genetics, high-throughput behavioral studies, and *in vivo* calcium imaging of chemosensory neurons (CSNs), we show that *C. elegans* can detect and respond to the drug, VPA, through a cyclic nucleotide-dependent signaling cascade. Wild-type animals are attracted to VPA in a manner that depends on the *tax-4* CNG ion channel and the *tax-4*-expressing AWC chemosensory neuron (CSN) pair. Furthermore, attraction to VPA is diminished in mutants for rGC-encoding genes, *odr-1* and *gcy-28*. As cGMP generators, *odr-1* and *gcy-28* may serve as direct VPA targets or act as downstream effectors used by signaling pathways that converge on *tax-4*. This uncertainty led us to explore the possibility that members of the GPCR family, which often signal through Gα subunits, contribute to VPA attraction. Assays with Gα mutants revealed a role for several genes encoding Gα proteins (*odr-3*, *egl-30*, and *gpa-2*;*gpa-3*) in mediating attraction to VPA, and consequently implicating GPCRs in the sensing of this drug. Thus, with its well-defined and highly sensitive nervous system, large repertoire of chemoreceptors, and conserved signaling components, *C. elegans* offers a unique genetic route to the analysis of VPA action.

## Methods

### Chemical and biological reagents

We purchased chemicals from the following commercial sources (supplier, catalog no.): valproic acid (APExBIO, B1251), furfural (TCI Chemicals, F00073), isoamyl alcohol (Sigma-Aldrich, W205710), 2-methyl-1-butanol (Sigma-Aldrich, W399817), and valeric acid (Sigma-Aldrich, 240370). We obtained *C. elegans* strains from our in-house collection, the *Caenorhabditis* Genetics Center, other members of the worm community, or derived new strains in-house. Supplementary Table 1 lists the genotypes and origin of all *C. elegans* strains used in this study.

### Growth and maintenance of *C. elegans* strains

We maintained and synchronized animals according to standard methods [28], using N2 (Bristol) as our designated wild-type (WT) and control strain. In brief, we grew all strains at 20°C on NGM plates seeded with OP50 *E. coli* as a food source and used well-fed young adult worms for all experiments. We maintained freshly thawed strains for at least three generations before conducting behavioral assays [29]. We discarded strains that had been maintained in the lab for more than 3-4 months. If needed for additional experiments, animals were thawed from frozen stocks.

### Calcium imaging lines

To measure calcium dynamics in *tax-4(p678), odr-1(n1396), or gcy-28(yum32)* mutants, we introduced transgenes driving expression of GCaMP6s in the nuclei of all CSNs (*aeaIs008[ift-20p::GCaMP6s::3x::NLS]* ?) and mCherry in the cytoplasm (*hpIs728[gpc-1p::mCherry]* X) of a subset of CSNs. Both transgenes were derived from ZM10104, a previously validated, CSN-specific calcium imaging line [30]. Except in the *odr-1* background, all strains carry the mCherry marker. We verified that lines contained *tax-4(p678)*, *odr-1(n1396)*, and *gcy-28(yum32)* by amplifying genomic DNA with gene-specific primers (see below) and sequencing the PCR-amplified product.

***tax-4* -** Forward: CCAATGGAATTGGCTCTCCTC, Reverse: CATCCCAAGTCAGGATACTG
***odr-1 -*** Forward: CGATAATTGGCCTTCTGCTC, Reverse: GGTCAACCTTCGTCCAATCCAC
***gcy-28 -*** Forward: agagaaacttttctaaacgtttttcaagaaaagc, Reverse: acttgaagaacggatcgtcctttaatcc

### Data acquisition for chemotaxis assays

We conducted multiple chemotaxis assays in parallel by adapting the workflow previously developed for high-throughput chemotaxis screening of natural products [21]. Briefly, we added 10 mL of chemotaxis media dissolved in molten Gelrite™ (Research Products International, 2.5% wt/vol) to each well of a 4-well multiwell plate (ThermoFisher, catalog no. 267061). Chemotaxis media contains (mM): potassium phosphate buffer (5) at ph 6; MgSO_4_ (1); and CaCl_2_ (1). To define the arena within each well, we used custom, 1mm-thick foam inserts cut by a computer-controlled cutting device (Cricut 3); inserts have an outer pentagon shape (border of the arena with a top-right notch) and an inner hexagon (the arena itself).

Unless otherwise noted, at one of the vertices of the hexagon, close to the notch, we added 2.5 μL of our compound of interest, and to the opposite vertex, 2.5 μL of solvent. Figure 1A illustrates the arrangement, including the craft foam insert design and the placement of the chosen compound and solvent vertices. We allowed our compounds to diffuse for 60 minutes prior to adding young adult animals to the assay arena. We washed well-fed, age-synchronized young adult *C. elegans* three times with chemotaxis buffer to separate animals from bacterial food. To initiate assays, we added a concentrated drop of worms to the start zone (Figure 1A) using a 200 μL single-channel micropipette and removed excess buffer with PVA eye spears (BVI, catalog no. NC0972725). To monitor the temperature and control relative humidity, we placed our chemotaxis assay plates inside a dehumidifying dry cabinet (Forspark, FSDCBLK30). We conducted our experiments at temperatures ranging from 21.9 °C to 25°C, with relative humidity levels ranging from 30% to 39%. We allowed the worms to explore the arena for 60 minutes. To capture the endpoint of the assay, we scanned the assay plates with a consumer-grade flatbed scanner (Epson Perfection V600, model: B11B198011). The focal plane of the scanner was adjusted to match the height of the chemotaxis media, maximizing image contrast [21]. We acquired images (1200 DPI/8-bit grayscale) and saved them as uncompressed *.tiff files. We used the following compounds and concentrations for the experiments in Figure 4C: furfural (20 mM), isoamyl alcohol (9.32 mM), and 2-methyl-1-butanol (20 mM), valeric acid (9.10 M), and valproic acid (6.12 M) [21].

**Figure 1:**
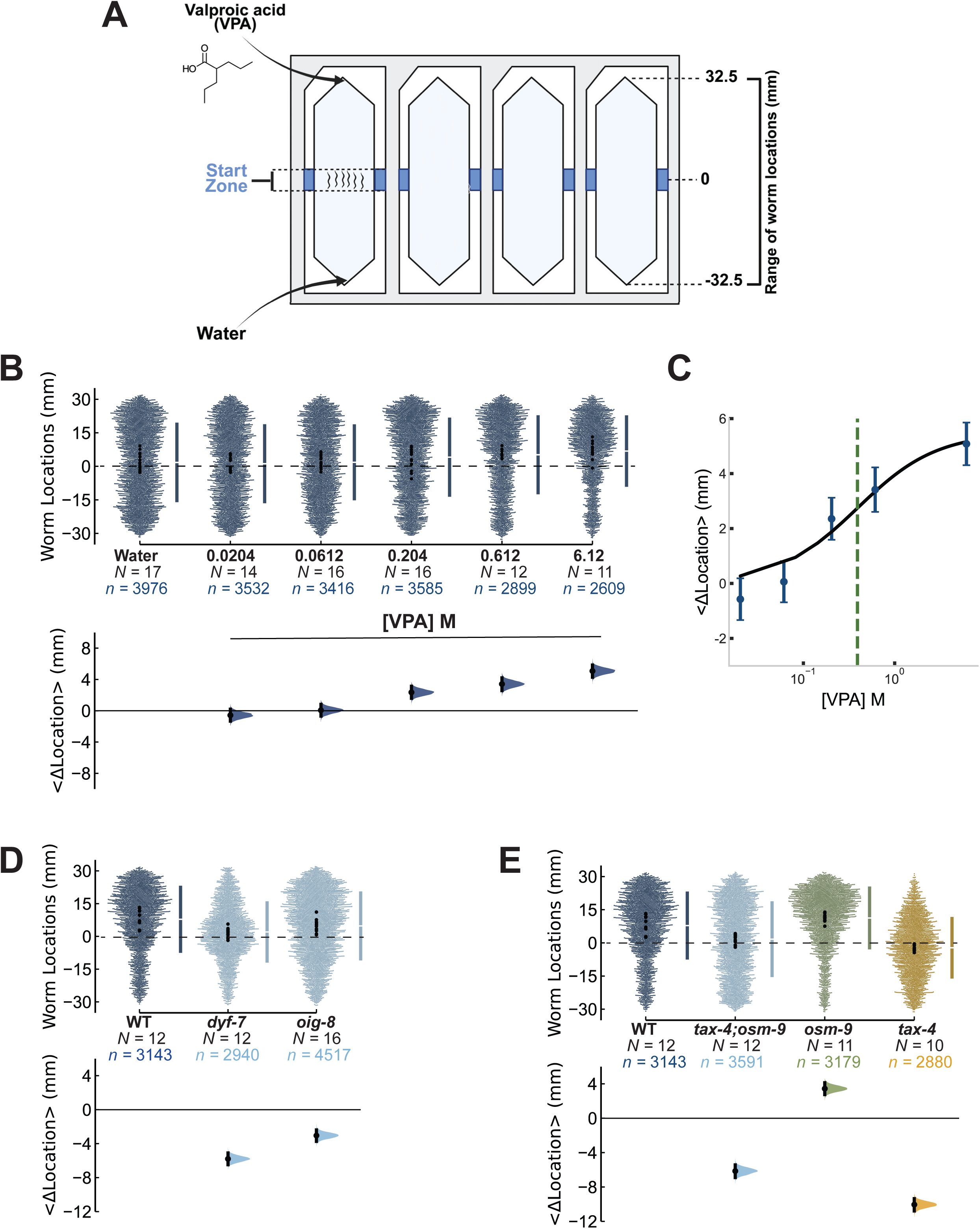
*C. elegans* are attracted to pure VPA in a *tax-4*-dependent manner. **(A)** Schematic of the 4-well chemotaxis assay plate. The locations (mm) for the start zone, VPA placement, and control (water) placement are 0, 32.5, and -32.5, respectively. Created in BioRender. Rogel-Hernandez, L. (2024). BioRender.com/y34o134. **(B)** Top: Swarm plot of the locations of individual animals (*n*, small dots) pooled across multiple assays (*N*, black dots). The mean location (gap) and standard deviation (represented by vertical lines) are shown to the right. Bottom: The bootstrapped mean difference relative to solvent controls (<Δlocation>) with its 95% confidence interval (CI). **(C)** Dose-response of VPA attraction, plotted as <Δlocation> ± 95% CI values from panel B *vs.* source concentration. The smooth line is a fit to data using a Hill equation, *EC*_50_ = 0.39 M, *n*_h_ = 1. **(D)** Defects in sensory cilia impairs attraction to pure VPA. Top: Small dots show the position of individual animals (*n*) pooled across multiple biological replicates (large dots, *N*). Bottom: Bootstrapped mean difference relative to WT controls with its 95% CI. **(E)** Loss of chemosensory ion channels impairs attraction to pure VPA. Top: Small dots show the position of individual animals (*n*) pooled across multiple biological replicates (large dots, *N*). Bottom: Bootstrapped mean difference relative to WT controls with its 95% CI.

**Figure 2:**
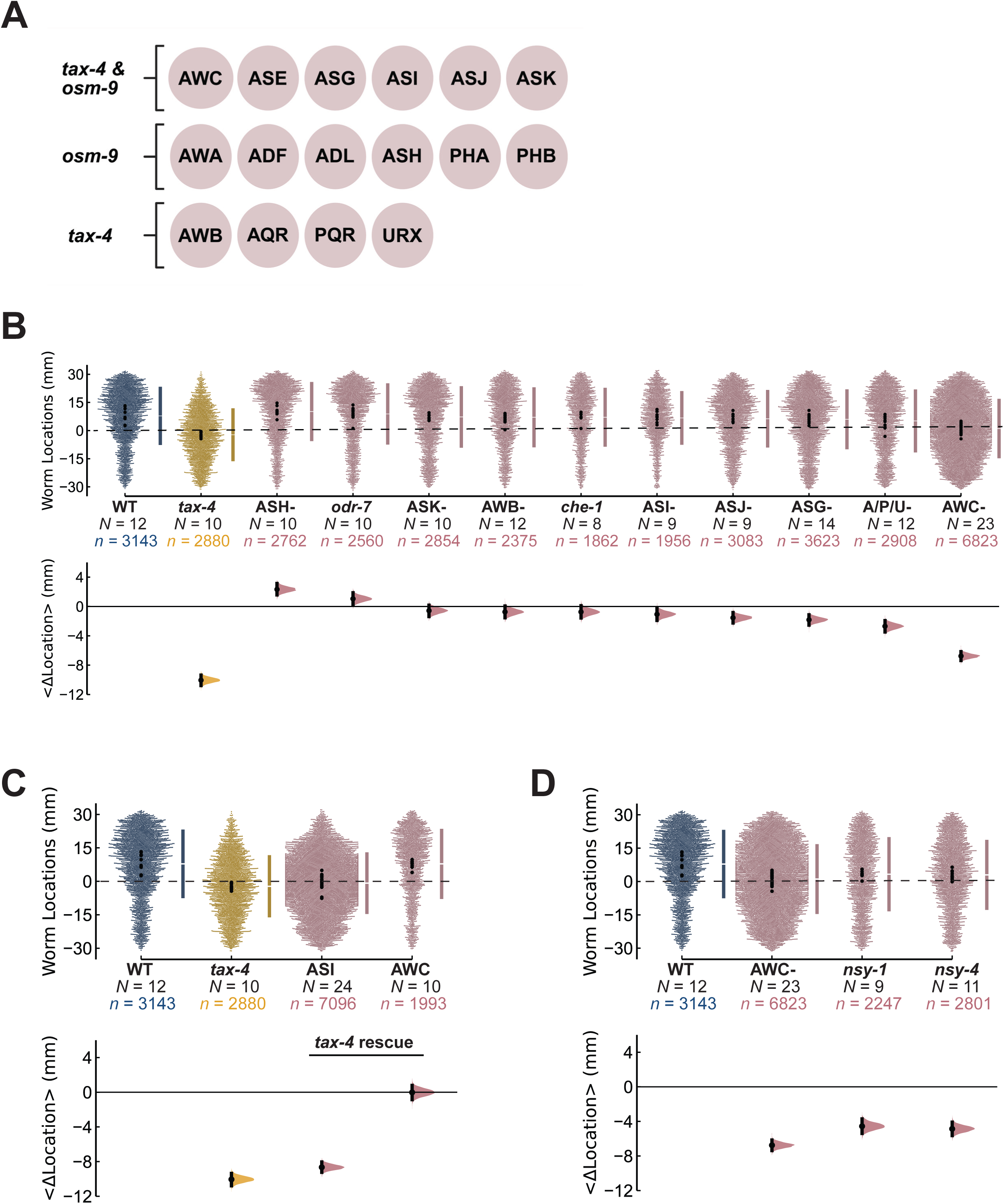
The *tax-4*-expressing AWC CSN pair mediates attraction to VPA. **(A)** Schematic showing expression of the *tax-4* and *osm-9* chemosensory ion channels in CSN pairs. Created in BioRender. Rogel-Hernandez, L. (2024) BioRender.com/v99y955. **(B-D)** VPA attraction as a function of loss of individual neurons **(B)**, neuron-specific rescue of *tax-4* expression **(C)**, and loss of AWC asymmetry **(D).** In each panel, the top row is a swarm plot of the locations of individual animals (*n*, small dots) pooled across multiple assays (*N*, black dots). The mean location (gap) and standard deviation (represented by vertical lines) are shown to the right. Bottom: Bootstrapped mean difference relative to WT controls with its 95% CI. For ease of visual comparison, the data for *tax-4* mutants in panel B and C are replotted from Figure 1E and the data for AWC- in Figure 2D are replotted from Figure 2B. Note, *nsy-1* mutant animals give rise to animals with two AWC^ON^ neurons while *nsy-4* to those with two AWC^OFF^ neurons.

### Data curation and analysis for chemotaxis assays

We assigned a unique identifier to all *.tiff images acquired and the assays represented in each image. We analyzed our images using Our Worm Locator (OWL) Graphical User Interface (GUI) to annotate the worm locations present in each assay [21]. Here, we define a worm location as a region within the arena occupied by at least one worm. For this study, assays with greater than 150 annotated worm locations were included in the analysis [21]. For each strain and test condition, we collected data in a blinded manner from at least three biological replicates. We aggregated the data for each strain and analyzed it using estimation statistics from the DABEST package (v. 0.3.1) [31]. We generated shared control estimation plots to compare all mutant strains (tests) against wild-type (control). When employing estimation statistics to analyze *C. elegans* responses to VPA, we can infer the following:

1. Positive <Δlocation> (aka bootstrapped mean difference) values whose 95% confidence interval (CI) does not intersect with the zero-intercept indicate that the mutant is, to an extent, more attracted to VPA than the control genotype.
2. For <Δlocation> values whose 95% CI falls at the zero-intercept, indicate that the mutant behavior is indistinguishable from the control genotype.
3. Negative <Δlocation> values whose 95% CI does not intersect with the zero-intercept correspond to the mutant having a partial or a complete loss of attraction to VPA compared to the control genotype.

In addition, we applied this statistical approach to compare the mean worm locations of animal populations exposed to different concentrations of VPA (tests) versus those exposed to water (control) (Figure 1B). Furthermore, to provide insights into how the <Δlocation> functions under different VPA concentrations, we decided to fit our data from Figure 1B to the Hill equation (Figure 1C), which is primarily used to describe dose-response relationships. Here, the Hill equation is represented in the following manner:

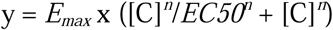

Where y represents the <Δlocation> at a concentration of VPA ([C]), *E_max_*is the maximum <Δlocation>, and *EC50* is the VPA concentration at which the <Δlocation> is half of the maximum, and *n* is the Hill coefficient, which was set to 1 to indicate a non-cooperative relationship. Supplementary Table 2 lists the mean worm location values for strains tested against VPA and other chemicals (refer to Figure 4C). Supplementary Table 3 lists the <Δlocation> values and their 95% CI quantified with the DABEST package for comparisons between control and test groups.

### Data acquisition for calcium imaging

We performed calcium imaging experiments, as previously described by Lin et al. (2023) [30]. To capture the calcium dynamics of 11 amphid CSN pairs, we immobilized individual transgenic *C. elegans* expressing the calcium indicator GCaMP6s in the nuclei of these neurons in a microfluidic device. To enable post-hoc neuronal identification, these animals also expressed cytoplasmic mCherry in the AFD, ASE, ASI, ASJ, AWB, and AWC neurons [30]. Transgenic animals retained the ability to migrate towards VPA in behavioral assays, underscoring their utility for determining cellular responses to VPA.

Unexpectedly, these transgenic animals were more strongly attracted to VPA than wild-type: <Δlocation> = 4.15 ± 95% CI [3.39, 4.88] mm. We exposed their noses to controlled pulses of VPA diluted with imaging buffer (mM): potassium phosphate (5, pH 6), MgSO_4_ (1), CaCl_2_ (1), and NaCl (50), adjusted to 350 mOsm with sorbitol (VPA final concentration: 6.12 x 10^-4^ M). Each animal underwent six 10-second pulses of VPA, separated by 30-second intervals of VPA-free imaging buffer. We repeated this procedure 2-4 times per animal to assess which amphid CSNs responded to VPA. To monitor the neuronal responses of *C. elegans* to VPA and similar to the work of Lin, et al [30], we used a spinning-disk confocal microscope equipped with a 60X objective to monitor GCaMP6s emission. We used a 488-nm laser to excite GCaMP6s and a 561-nm laser to excite mCherry. We captured volumetric data at 2.5 frames per second (fps)1.6 fps (*gcy-28*), or 1.2 fps. We imaged control and *odr-1* animals at 2.5 fps, *gcy-28* animals at 1.6 fps, and 62% of *tax-4* animals at 2.5 fps and 38% at 1.2 fps.

### Data curation and analysis for calcium imaging

We assigned unique identifiers to each animal and its corresponding recordings to manage and process the raw data. We obtained recordings across four strains: control (ZM10104), *tax-4* (GN1103), *odr-1* (GN1126), and *gcy-28* (GN1136). The number of animals recorded from each strain is as follows: control (*N* = 21 animals), *tax-4* (*N* = 21), *odr-1* (*N* = 7), and *gcy-28* (*N* = 5). From these animals, we collected a total of 155 recordings: control (*N =* 59 recordings), *tax-4* (*N =* 61), *odr-1* (*N =* 21), and *gcy-28* (*N =* 14). Because the frame rate we used to acquire image stacks varied between 1.2 and 2.5 fps (see above), we interpolated all datasets to 2.5 fps (control, *tax-4*, *odr-1*) or to 1.6 fps (*gcy-28*) prior to analysis. We used MATLAB to perform all image processing tasks, which included identifying and tracking CSNs, extracting their corresponding calcium signals, normalizing these signals, and averaging them per CSN pair across animals [30]. Refer to Lin et al. (2023) [30] for details about how we identified, tracked, and analyzed individual neurons. Briefly, for each recording, the orientation of each animal was determined to properly locate the positions of all anterior CSNs. Given that ASH and ASJ are easily identified, we annotated these two pairs first, followed by three pairs that are close to each other: ASK, ADL, and ASI [30]. We annotated the rest of the cells with the guidance of the cytoplasmic mCherry marker and with respect to their position relative to other identifiable neurons [30]. An experienced observer assigned each neuron its identity based on the well-defined neuroanatomy of the *C. elegans* nervous system [32] and the position of each neuron relative to the body, feeding organ (pharynx), and to mCherry-expressing neurons. We would like to note, however, that the absence of the mCherry marker in the *odr-1* background and the reduced sensitivity to VPA in this mutant background complicated CSN annotation. Other instances in which it was difficult to annotate the identity of CSNs occurred when the neurons were too close to each other or lacked detectable stimulus-linked in GCaMP6s signal. To track the CSNs over time, we made use of a neighborhood correlation tracking algorithm, where the center of each neuron and its surrounding 3D neighborhood are tracked independently from frame to frame [30]. The baseline fluorescence was determined individually for each recorded neuron. Fluorescence, *F*, was monitored over time and corrected for photobleaching; we reported signals as Δ*F/F*_0_, with *F*_0_ defined as the fluorescence recorded prior to VPA stimulation. We proofread the resulting traces and excluded any that (a) showed signal mixing with neighboring neurons (we confirmed this by verifying neuron overlap in the recorded images); (b) showed signs of mistracking (e.g. sudden signs in activity caused by the tracked point jumping to a different nearby neuron); or (c) showed large changes in baseline activity during the recording (sometimes caused by motion artifacts when the head of the worm drifts over time) that prevented us from establishing a consistent *F*_0_.

To compare the different calcium imaging lines to the control during VPA presentation and assess the drug’s effect, we computed the mean global signed max Δ*F/F*_0_ value (Supplementary Table 4) for each CSN pair for each calcium imaging line and scored statistical significance by using the Mann-Whitney U test (Supplementary Table 5). As a first step, we applied a Savitzky-Golay filter with third order polynomial to each recording to smooth the signals and reduce noise. We then proceeded to assess the magnitude of the response for each CSN pair to VPA by identifying both their global maximum and minimum values within the time window in which the drug was applied, and the one with the larger absolute value was selected as the signed max ΔF/F_0_ value. We employed this single-point metric to accommodate the natural variability in responses across all eleven pairs of CSNs to VPA.

## Data and Code Availability

The code used to convert images of chemotaxis assay plates into animal position, OWL, is freely available on GitHub: https://github.com/Neuroplant-Resources/Neuroplant-OWL.git via a digital object identifier, DOI: https://doi.org/10.5281/zenodo.7807007.

## Results

### *C. elegans* is attracted to VPA in a dose- and *tax-4*-dependent manner

Although it is well-established that VPA has valuable therapeutic benefits for the treatment of epilepsy and bipolar disorder, the chemical signaling pathways engaged by VPA in the nervous system remain enigmatic. To address this knowledge gap, we took an unconventional approach and explored the possibility that VPA induces chemoattraction or repulsion in *C. elegans*. Using a high-throughput chemotaxis-based platform [21], we found that wild-type (WT) animals are attracted to VPA and that the strength of attraction increases with the source concentration (Figure 1B). The *EC*_50_ for attraction is 0.39 M at the source location. To evoke the strongest possible behavioral response, we delivered pure VPA (6.12 M) at the source position for all subsequent behavioral tests. Notably, animals are not immersed in VPA during these assays, minimizing the risk of toxic effects. Our finding that animals are able to migrate toward the VPA source implies that they can sense VPA at the starting zone, where the concentration is likely to be several orders of magnitude less than that at the source.

Next, we sought to evaluate the role of the chemosensory nervous system in mediating this behavior. Eleven pairs of ciliated CSNs reside in the amphid sensilla in the head and five in the phasmid sensilla in the tail [22]. Although these CSNs possess distinct sensory functions, they are all bipolar sensory neurons bearing ciliated sensory dendrites and an axon emanating from the cell body [22,33].

Thus, to assess if one or multiple CSN pairs detect VPA, we tested *dyf-7* and *oig-8* mutants with defects in sensory dendrites or cilia. Without functional *dyf-7*, a subset of head CSNs fail to extend their dendrites to the tip of the nose, while the cilia and axons develop normally [33]. This morphological defect displaces the cilia and its chemoreceptors further away from the nose opening, thereby reducing their exposure to several environmental chemical cues and ultimately impairing their ability to detect them [33]. Meanwhile, *oig-8* mutants give rise to animals with altered ciliary branching in AWA, AWB, and AWC CSNs [34].

In surveying the ability of *dyf-7* and *oig-8* mutants to sense VPA via the use of chemotaxis assays, we found that both mutants exhibit a reduction in attraction to this drug in comparison to wild-type animals, as indicated by a negative <Δlocation> value. This finding implicates cilia in the first steps of VPA sensing. Next, we tested mutants in the *tax-4* (CNG) and *osm-9* (TRP) ion channel genes required for chemosensory transduction [22]. We found that *tax-4;osm-9* double mutants are indifferent to VPA (Figure 1E), confirming the central role for CSNs and chemosensory signaling in VPA sensing. Since these ion channels are differentially expressed within the chemosensory nervous system, we tested single mutants for these two channels to narrow VPA’s site of action [21,35]. We found that *osm-9* single mutants, but not *tax-4* mutants, were attracted to the drug (Figure 1E). Unexpectedly, VPA attraction was modestly enhanced in *osm-9* single mutants (Figure 1E), suggesting that, similar to attraction to isoamyl alcohol and 2-methyl-1-butanol [36]*, osm-9* functions as a negative regulator of VPA attraction. Together, these findings strongly implicate sensory cilia and one or more *tax-4-*expressing CSN as candidate primary receptor neurons for VPA.

### VPA attraction relies on the *tax-4*-expressing AWC chemosensory neuron pair

Having established that *C. elegans* is attracted to VPA in a *tax-4-*dependent manner, we next sought to determine which *tax-4*-expressing neurons are required for this behavior. To achieve this, we collected mutants overexpressing constitutively active caspases under cell-specific promoters, a strategy that induces apoptosis and eliminates the target cell [37]. For nine out of the ten *tax-4*-expressing CSNs (Figure 2A), we tested the ability of mutants lacking individual CSN pairs (AWB, AWC, ASG, ASI, ASJ, and ASK) or in combination (AQR, PQR, and URX) to detect VPA. To assess whether ASE plays a role in VPA sensing, we tested mutants lacking the *che-1* transcription factor, which is required for ASE identity and function [38]. In addition to assessing the roles of these *tax-4*-expressing CSNs in detecting VPA, we also tested animals lacking two *osm-9*-expressing CSNs, ASH and AWA, with the expectation that they might behave similarly to *osm-9* mutants. For ASH, we used a line expressing constitutively active caspases under an ASH-specific promoter [39]. For AWA, we assessed the ability of *odr-7* mutants to respond to VPA, as this mutant fails to detect AWA-sensed chemicals and is reported to adopt an AWC-like fate [40].

In comparing the behavior of these mutants to those of intact, wild-type animals, it is clear that animals lacking the *tax-4* expressing CSN pair AWC contribute significantly to the detection of VPA (<Δlocation> = -6.76 ± 95% CI [-7.37, -6.16] mm) and essentially phenocopy the *tax-4* phenotype (Figure 2B). The remaining *tax-4* expressing CSNs contribute moderately (AQR, PQR, URX, abbreviation: A/P/U-), minimally (ASJ, ASG), or not at all (ASK, AWB, ASE, and ASI) to VPA attraction (Figure 2B). As proposed above, animals missing the *osm-9*-expressing ASH CSN pair are slightly more attracted to VPA (<Δlocation> = 2.33 ± 95% CI [1.58, 3.14] mm) than wild-type and display similar responses as those observed for *osm-9* single mutants (Figure 1E, <Δlocation> = 3.44 ± 95% CI [2.75, 4.13] mm). On the other hand, *odr-7* mutants with disrupted AWA neuron fate behave like wild-type and remain attracted to VPA. These findings demonstrate that AWC neurons are required for VPA attraction and leave open the possibility that additional CSNs may contribute to this behavior.

To determine whether or not *tax-4* is required in the AWC neurons, we restored wild-type *tax-4* expression to the AWC in a *tax-4* null background (Figure 2C). We found that wild-type *tax-4* expression in AWC alone was sufficient to re-establish wild-type VPA attraction, indicating that *tax-4* acts cell-autonomously within the AWC neurons to promote VPA attraction. By contrast, restoring wild-type *tax-4* to ASI, which provides synaptic input to AWC [32], had no effect on VPA attraction (Figure 2C). Unlike most other CSN pairs, the AWC pair exhibits asymmetric expression of chemoreceptors [41]. The AWC^ON^ neuron expresses the GPCR *str-2*, while the AWC^OFF^ neuron lacks *str-2* expression and expresses the GPCR *srsx-3* [41]. Each worm contains one AWC^ON^ and one AWC^OFF^ neuron [41]. To investigate whether one or both AWCs mediate attraction to VPA, we conducted chemotaxis assays with mutants that give rise to either two AWC^ON^ (*nsy-1*) or two AWC^OFF^ (*nsy-4*) neurons [41]. Like animals lacking AWC neurons entirely, *nsy-1* mutants lacking AWC^ON^ neurons and *nsy-4* mutants lacking AWC^OFF^ neurons are not attracted to VPA (Figure 2D), suggesting either that both types of AWC neuron contribute to VPA-evoked chemotaxis or that some factor conferred by asymmetry in AWC signaling is needed for this behavior. These findings reinforce two ideas, namely that *tax-4* acts within the AWC neurons and this pair of CSNs are essential for VPA attraction.

### VPA evokes *tax-4*-dependent calcium transients in AWC, AWB, and ASH

To better understand the contribution of individual CSNs in VPA-sensing in an intact animal, we imaged VPA-evoked changes in intracellular calcium in 11 amphid CSNs [30], superfusing the animal’s nose with VPA diluted to a final concentration of 0.6 mM in physiological saline. This concentration mirrors the drug levels (0.35-0.69 mM) found in the plasma of VPA-treated patients [42]. In control animals, the AWC and AWB CSNs stood out as VPA-sensitive (Figure 3, Supplemental Figure 1A). These experiments revealed that the AWC neurons act as the primary cellular sensors for VPA, agreeing with our results from the chemotaxis assays with lines missing different CSN pairs. The AWC neurons responded consistently and robustly to VPA, evoking large negative calcium transients upon its presentation and large positive calcium transients upon its removal (Figure 3). These so-called OFF responses are a hallmark of AWC chemosensory transduction [22]. Interestingly, the AWB neurons responded to VPA presentation in an ON manner in roughly one out of every four animals (Supplementary Figure 1A). The variability observed in AWB could stem from differences in gene expression among animals or that the AWB neurons are less sensitive to VPA and would exhibit stronger responses to higher VPA doses. These findings underscore the central role of the AWC neuron pair in VPA sensing and establish that, similar to neurons in other animals, including mammals [7–16], *C. elegans* neurons are sensitive to submillimolar concentrations of VPA.

**Figure 3:**
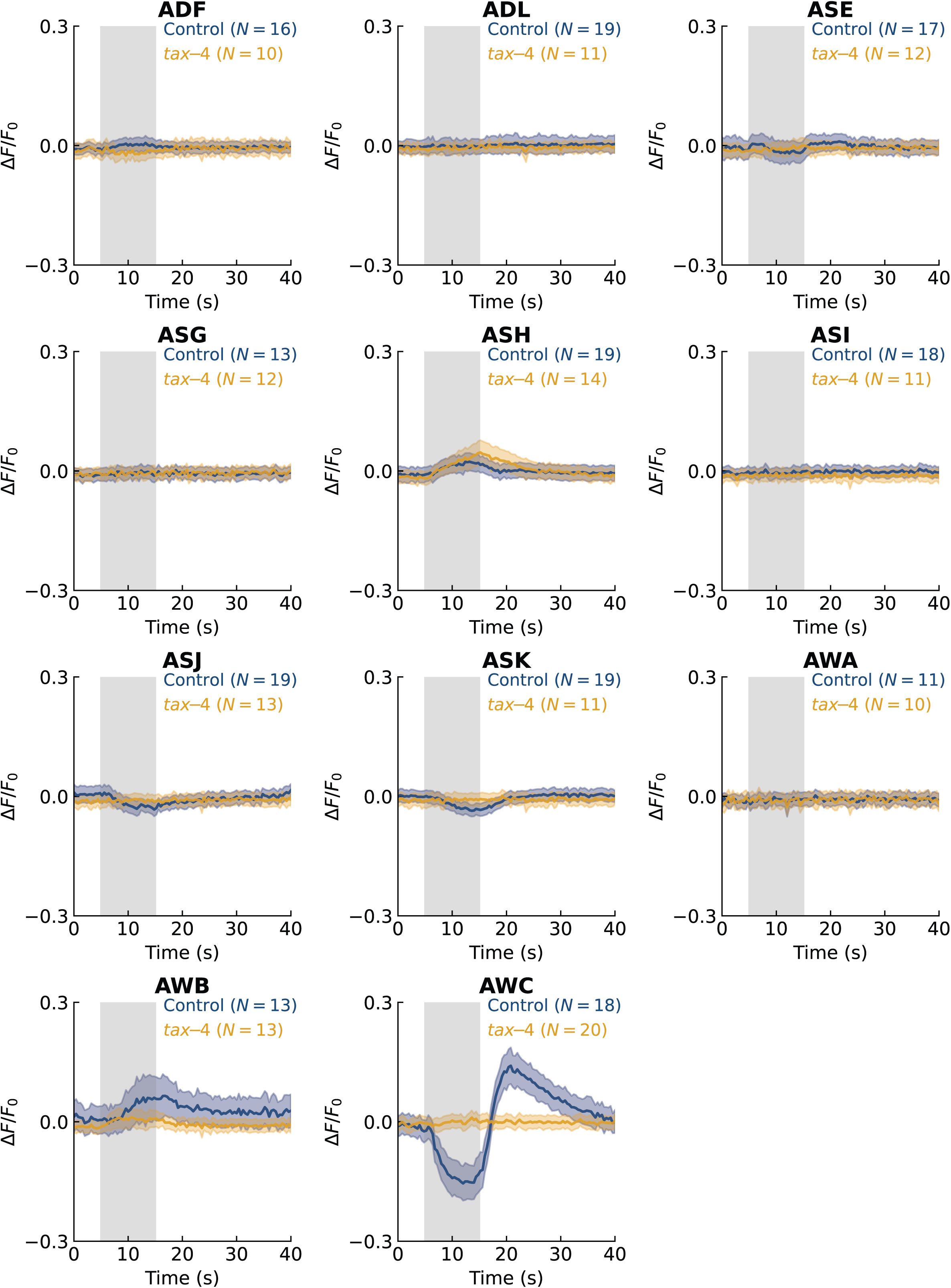
VPA evokes robust *tax-4*-dependent OFF responses in AWC and variable ON responses in AWB and ASH. The neuronal activity of all amphid CSNs from control (navy blue) and *tax-4* (golden yellow) calcium imaging lines in response to VPA (6.12 × 10^-4^ M, 1:10^-4^ dilution) presentation and removal averaged across multiple trials (*N*). A shaded gray box indicates the 10-second VPA delivery period. Navy blue and golden yellow shaded areas represent the standard error of the mean (SEM) for control and *tax-4*, respectively.

**Figure 4:**
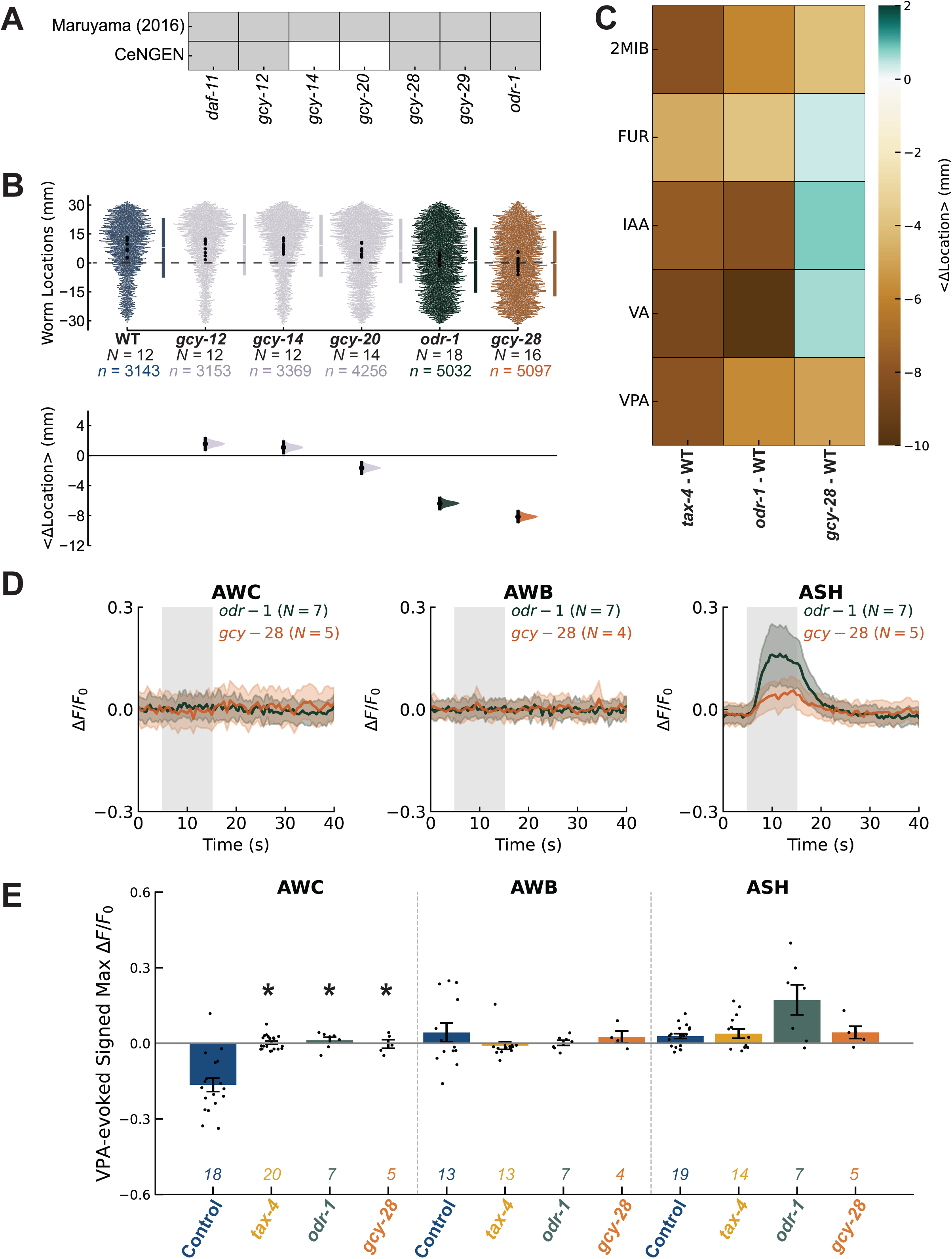
Attraction to VPA is mediated by two rGCs, *odr-1* and *gcy-28*. **(A)** Schematic of the expression of candidate rGC genes for the AWC CSN pair, based on Maruyama (2016) and adult CeNGEN single-cell RNA-seq dataset [24,46,47]. Gray shading indicates the presence of rGC genes. **(B)** Comparison of five rGC mutants to WT animals in VPA attraction assays. Top: Swarm plot of the locations of individual animals (*n*, small dots) pooled across multiple assays (*N*, black dots). The mean location (gap) and standard deviation (represented by vertical lines) are shown to the right. Bottom: Bootstrapped mean difference relative to WT controls with its 95% CI. **(C)** Heatmap of <Δlocation> values depicting pairwise comparisons between *tax-4*, *odr-1*, and *gcy-28* to WT in response to five compounds: isoamyl alcohol (IAA), furfural (FUR), valeric acid (VA), 2-methyl-1-butanol (2M1B), and valproic acid (VPA). Columns represent genotype comparisons, and rows represent the different attractants tested. Shades of green are positive <Δlocation> values, while shades of brown represent negative <Δlocation> values. The deeper the shade of green or brown, the greater the magnitude of the <Δlocation> value and the greater the deviation from WT behavior. **(D)** The neuronal activity of AWC, AWB, and ASH from *odr-1* (dark green) and *gcy-28* (burnt orange) calcium imaging lines in response to VPA (6.12 × 10^-4^ M, 1:10^-4^ dilution) presentation and removal averaged across multiple trials (*N*). A shaded gray box indicates the 10-second VPA delivery period. Dark green and burnt orange shaded areas represent the SEM for *odr-1* and *gcy-28*, respectively. **(E)** Quantification of the signed max ΔF/F_0_ value during VPA presentation for AWC, AWB, and ASH across four calcium imaging lines: control, *tax-4*, *odr-1*, and *gcy-28*. Bars represent SEM and italicized numbers above tick marks in x-axis indicate the number of trials (*N*) per each CSN pair. Statistically significant comparisons (*p* < 0.05) as a function of genotype in each neuron are annotated with an asterisk (Mann-Whitney U test).

To determine how the *tax-4* CNG channel contributes to VPA sensing by CSNs, we recorded the activity of amphid CSNs in a *tax-4* null background (Figure 3, Supplemental Figure 1B). Consistent with a central role for *tax-4* in VPA-evoked signaling, loss of *tax-4* abolished VPA-evoked calcium transients in AWC and AWB. We also observed *tax-4*-dependent VPA-evoked responses in ADF, ASE, ASH, ASJ, and ASK (Figure 3, Supplementary Figure 1), suggesting that these neurons might also contribute to VPA sensing. Unexpectedly, loss of *tax-4* increased the frequency (7/14 vs. 8/19 of trials in *tax-4* and control animals) and amplitude of VPA-evoked calcium transients in the ASH neurons (Figure 3, Supplementary Figure 1). Because the ASH neurons do not express *tax-4*, these findings imply that a *tax-4*-expressing CSN modulates ASH signaling during VPA application. Together, these findings reveal that 7 of 11 amphid CSN pairs exhibit *tax-4*-dependent, VPA-evoked calcium transients and that the AWC neurons have the strongest, most robust responses.

It has long been known that the ASE and AWC neurons exhibit left-right asymmetry in the expression of membrane receptors (reviewed in Ref. [43]). To explore whether or not sensitivity to VPA is asymmetric, we analyzed VPA-evoked calcium transients separately in the left and right neurons (Supplemental Figure 2). We found that ASER, but not ASEL, is sensitive to VPA even though both neurons express the *tax-4* ion channel gene. This directional asymmetry suggests that only the ASER neurons express VPA-sensitive receptors. By contrast, both the left and right AWCs responded to VPA in a *tax-4-*dependent manner (Supplemental Figure 2) Because asymmetric receptor expression is determined stochastically in the AWC neurons [44], we are not able to link responses in the left and right AWC neurons to their asymmetric identity as either AWC^ON^ or AWC^OFF^ neurons. Considered together, *in vivo* calcium imaging affirms the central role of the AWC neurons in VPA sensing and implicates ASER, AWB, and ASH as additional signaling partners.

### Two receptor guanylyl cyclase genes, *odr-1* and *gcy-28*, are required for VPA attraction

Given the dominant role of AWC in both cellular and behavioral responses to VPA and its dependence on expression of the *tax-4* CNG ion channel gene, we next sought to identify the receptor guanylyl cyclases (rGCs) involved in VPA attraction. This class of membrane proteins plays a conserved role in CNG channel activity in sensory neurons [45]. We initiated our exploration by analyzing the VPA-evoked behavior of mutants lacking five rGC genes known to be expressed in AWC (Figure 4A; Refs. [24,46,47]). We found that *gcy-14, gcy-12,* and *gcy-20* mutants performed similarly to wild-type or exhibited minor defects (Figure 4B) and that *odr-1* and *gcy-28* mutants were indifferent to the drug (Figure 4B). Thus, only a subset of the AWC-expressed rGC genes are required for VPA sensing.

Because *odr-1* and *gcy-28* have been previously implicated in sensing other chemical stimuli [22,48–50], it seems unlikely that either rGC is specifically required for VPA sensing. We directly tested this inference by assaying *odr-1* and *gcy-28* mutants for their ability to detect four additional compounds: furfural (FUR), isoamyl alcohol (IAA), 2-methyl-1-butanol (2M1B), and valeric acid (VA), an analog of VPA [1]. We selected this panel of compounds because, like VPA, they all attract *C. elegans* in a *tax-4*-dependent manner [21,36,51]. As expected from previous findings, attraction to each of these compounds requires *tax-4* signaling pathways (Figure 4C). In addition to mediating attraction behaviors to VPA and IAA, *odr-1* is essential for attraction to FUR, 2M1B, and VA (Figure 4C). On the other hand, *gcy-28* is crucial for mediating attraction to VPA and 2M1B, as its absence leads to a reduction in sensitivity to these compounds (Figure 4C). Interestingly, *gcy-28* mutant animals appear slightly more attracted to FUR, IAA, and VA relative to wild-type (Figure 4C). Thus, *odr-1* in is required to detect our entire panel of attractants and *gcy-28* is required for attraction to 2MIB and VPA. From these findings, we infer that neither rGC is a candidate VPA receptor and that one or both function as downstream effectors of VPA receptors.

To assess whether *odr-1* and *gcy-28* are essential for facilitating VPA sensing in AWC, we monitored the calcium transients of this CSN pair in response to VPA in mutant backgrounds (Figure 4D). These experiments revealed that in the absence of either rGC, the calcium transients observed in control animals for AWC during VPA presentation are absent, similar to the *tax-4* mutant background (Figure 4E). This was also the case for AWCL and AWCR (Supplementary Figure 3), suggesting that *odr-1* and *gcy-28* are tightly linked to VPA sensing in both AWC^ON^ and AWC^OFF^. None of our AWC recordings in a *gcy-28(yum32)* [52] displayed detectable VPA-evoked calcium transients (Supplementary Figure 3), which differs from a prior study that found no defect in butanone-, isoamyl alcohol- and benzaldehyde-evoked AWC^ON^ calcium signals in *gcy-28(tm2411)* [50]. This discrepancy could indicate that *gcy-28* is required for AWC chemosensory responses to VPA, but not for responses to other chemical stimuli. VPA has a very low vapor pressure compared to the other chemicals, suggesting a role for *gcy-28* in generating responses in AWC to non-volatile chemicals like VPA. Alternatively, the loss of chemical sensitivity in *gcy-28* mutants in this study could indicate that the molecular lesions induced by the *yum32* allele generate a stronger phenotype than the *tm2411* allele. Additional experiments will be needed to resolve this discrepancy. Building on our finding that *odr-1* and *gcy-28* mutants are both defective in VPA-sensing by the AWC neurons and prior work showing that ODR-1 is concentrated in the cilia [48], while GCY-28 is confined to the axons [50], it is tempting to postulate that these two rGCs work together in a two-step signaling pathway to relay VPA detection and activate the AWC neurons with cGMP acting as the key secondary messenger in both locations [50].

In addition to AWC, we captured the calcium transients of the remaining amphid CSN pairs to VPA in *odr-1* and *gcy-28* mutant backgrounds (Supplementary Figure 4). Similar to AWC, the calcium transients previously observed in AWB are absent in *odr-1* and *gcy-28* (Figure 4D, 4E). However, given AWB’s variability in VPA-evoked calcium transients, we cannot exclude the possibility that these mutants do not affect VPA-evoked calcium transients in AWB (Figure 4E). Interestingly, the ASH neurons, much like in control and *tax-4* backgrounds, still respond to VPA, and in a more pronounced manner in the *odr-1* background (Figure 4D, 4E). The small and variable calcium transients observed for ADF are mediated by *gcy-28,* while those for ASK, ASJ, and ASE are not mediated by either rGC (Supplementary Figure 5). Given that ASE is an asymmetric pair, and in control animals, most responses to VPA are mediated by ASER, we analyzed calcium transients of the left and right ASE neurons independently from one another to assess if averaging these pairs together obscured valuable insights (Supplementary Figure 3). Similar to control animals, ASER responded to VPA in a subset of recordings in both mutant backgrounds and thus, neither rGC is essential for facilitating this cell’s ability to detect VPA (Supplementary Figure 3). Since ASER activity relies on *tax-4*, it is very likely that another rGC is involved in the detection process that may or may not be a direct target for VPA.

We identified an important role for *odr-1* and *gcy-28* in VPA-evoked calcium transients in the AWC, AWB, and ASH neurons, but not in the ASE neurons. These findings inform a model where attraction to VPA engages multiple CSNs to induce a certain level of attraction and this behavior is strongly driven by cGMP-dependent signals with overlapping, but non-identical molecular signaling pathways. The central role of the AWC neurons in VPA attraction further supports a model in which these neurons detect VPA in a manner that depends on the ODR-1 and GCY-28 rGCs and the TAX-4 ion channel proteins and that none of these three proteins confer selective sensitivity to VPA or bind VPA.

### Disruption of G protein signaling impairs VPA attraction

What other class of chemoreceptor proteins might grant the AWC neurons the ability to detect the presence of VPA? G-protein coupled receptors (GPCRs) are cell surface receptors that facilitate physiological responses to endogenous and environmental chemical stimuli, including odorants and tastants [27]. Similar to the model emerging from this study for VPA-evoked signaling in AWC, signaling pathways that mediate vertebrate vision depend on GPCRs that converge on CNG channels and include G proteins and rGCs as intermediate effectors [53]. Whereas the *C. elegans* genome contains more than 1500 genes predicted to encode GPCRs [25], there are only 21 genes encoding Gα subunits [54], 1 gene encoding β-arrestin [55], and 13 genes encoding regulators of G protein signaling (RGS) [56]. As illustrated in Figure 5A, all of these genes encode for proteins that are central to classical GPCR signaling [57]. First, we obtained mutants defective in the function of 8 (of 10) genes encoding Gα proteins expressed in the AWC neurons (Figure 5B) and tested them in VPA chemotaxis assays. In this way, we identified at least four Gα genes required for VPA attraction (Figure 5C). The *odr-3*, *egl-30*, and the double mutant *gpa-2;gpa-3* displayed the most severe defects in VPA attraction (Figure 5C). Since our *gpa-2;gpa-3* double mutant exhibited a strong defect, we also tested *gpa-2* and *gpa-3* single mutants. Interestingly, *gpa-2* mutants show a slight increase in attraction to VPA compared to WT, whereas *gpa-3* mutants exhibit a decrease in attraction to VPA. These results indicate that *gpa-2* and *gpa-3* have distinct roles in the neural circuit pathway responsible for mediating VPA attraction, and the absence of both disrupts the pathway’s function more severely than either mutation alone. The *odr-3* mutant was significantly impaired in detecting VPA (Figure 5C). Notably, *odr-3* expression is limited to the chemosensory nervous system. Although *odr-3* mutations are known to disrupt the morphology of AWC cilia [58], such morphological defects do not necessarily impair AWC signaling [59]. Thus, *odr-3* is a strong candidate GL gene for mediating VPA sensing. The other Gα mutant with a large effect, *egl-30*, also implicates a GPCR in mediating attraction to VPA (Figure 5C). However, because *egl-30* mutants exhibit a sluggish phenotype, we cannot distinguish between defects due to changes in movement or to a loss of VPA sensing. The remaining Gα mutants tested were more similar to wild-type (Figure 5C), suggesting that they have little or no influence on the overall response to VPA. Although further research is needed to determine which Gα subunits contribute to VPA-sensing by the AWC neurons, these results establish a role for G proteins and potentially GPCRs in VPA sensing.

**Figure 5:**
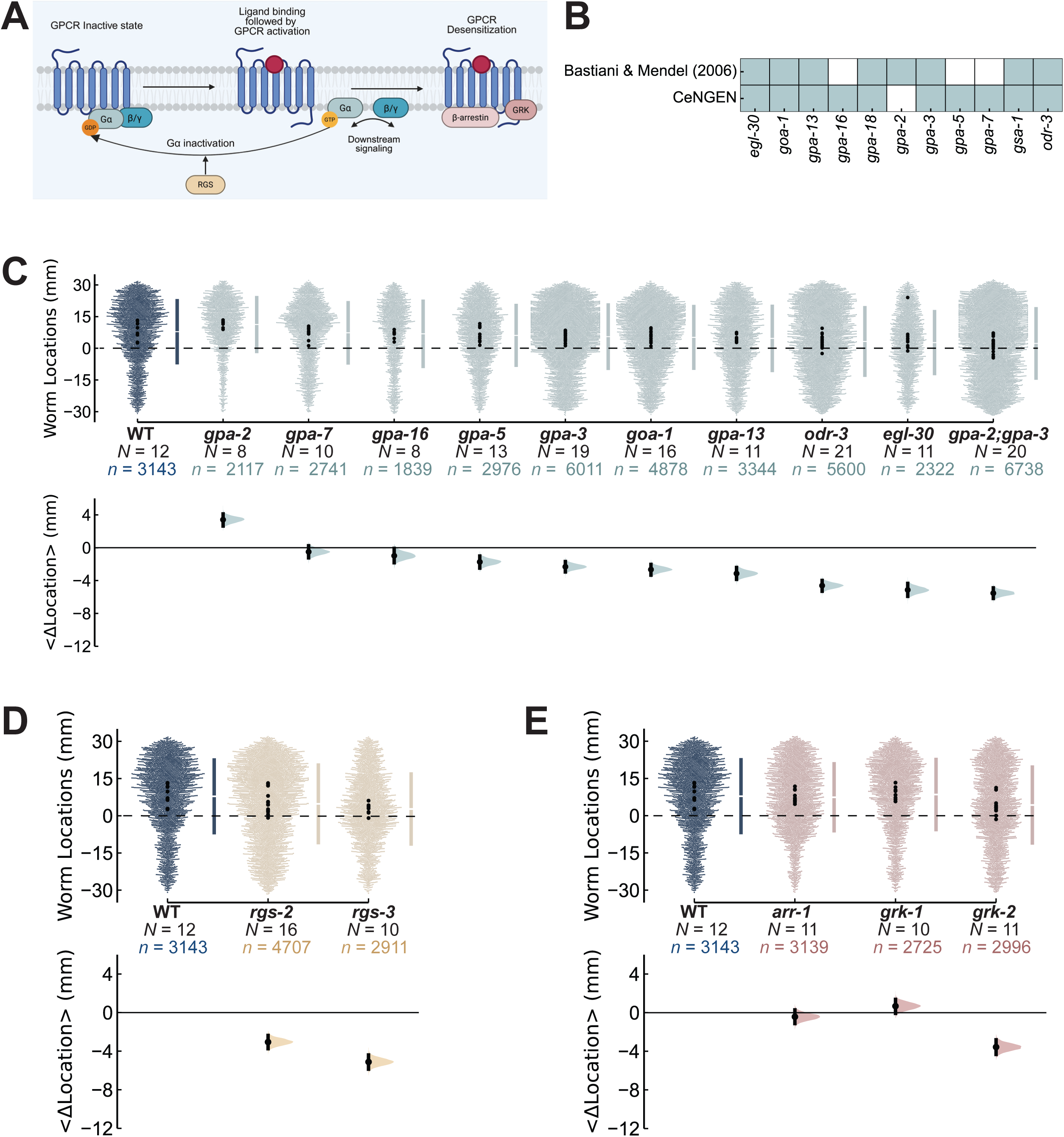
VPA sensing requires G proteins and GPCR signaling partners. **(A)** Schematic depicting the mechanisms by which G protein subunits, RGS, β-arrestin, and GRK proteins interact with GPCRs. In its inactive state, a GPCR is bound by a heterotrimeric G protein complex. Upon ligand binding, the GPCR acts as a GEF, exchanging GDP-bound Gα for GTP, thereby dissociating the heterotrimeric G protein into two active subcomplexes (Gα and Gβγ), which recruit downstream effectors to amplify GPCR signals. RGS proteins facilitate the reversion of the Gα to the GDP-bound state. To desensitize the GPCR, GRK proteins phosphorylate the C-terminal tail of the receptor, recruiting β-arrestin and promoting its binding to the receptor. This series of events prevents the receptor from activating G proteins and its targeted for endocytosis. Created in BioRender. Rogel-Hernandez, L. (2025) https://BioRender.com/dqe4xco. **(B)** Schematic of 11 candidate Gα subunits expressed in the AWC CSN pair, based on Bastiani and Mendel (2006) and adult CeNGEN single-cell RNA-seq dataset [46,47,54]. Light blue shading indicates the presence of the Gα gene. **(C-E)** Top: Swarm plot of the locations of individual animals (*n*, small dots) pooled across multiple assays (*N*, black dots). The mean location (gap) and standard deviation (represented by vertical lines) are shown to the right. Bottom: Bootstrapped mean difference relative to WT controls with its 95% CI. Panel **(C)** compares the responses of Gα mutants to those of WT. Panel **(D)** compares the responses of *rgs-2* and *rgs-3* mutants to those of WT. Panel **(E)** compares the responses of *arr-1*, *grk-1*, and *grk-2* mutants to those of WT.

To further evaluate the role of GPCR signaling in AWC, we conducted chemotaxis assays with *rgs-2* and *rgs-3* mutants (Figure 5D). These genes encode RGS proteins, which function to enhance the activity of Gα proteins, promoting their transition to the inactive GDP-bound state (Figure 5A) [56]. The gene *rgs-2* is expressed broadly within the worm, including in the AWC neurons [60]. On the other hand, *rgs-3* is exclusively expressed in nine pairs of CSNs, including the AWC neurons [61]. Without *rgs-2* or *rgs-3*, the animals no longer respond to VPA to the same extent as wild-type (Figure 5D). These findings are consistent with prior reports indicating that loss of *rgs-3* increases GPCR signaling and reduces sensory responses by decreasing synaptic transmission [22,61].

Finally, we surveyed the ability of *arr-1*, *grk-1*, and *grk-2* mutants to detect VPA (Figure 5E). The gene *arr-1* encodes for arrestin, while *grk-1* and *grk-2* encode for two distinct GPCR kinases (GRKs). Among other functions, GRKs and arrestin work together to desensitize GPCRs and protect cells from receptor overstimulation (Figure 5A) [57]. Signaling through many GPCRs via G proteins typically ends with the phosphorylation of the active receptor by GRKs, followed by arrestin binding and receptor internalization [57]. Our chemotaxis assays revealed that *arr-1* and *grk-1* mutants behave like wild-type and that *grk-2* mutants were defective in VPA attraction. These findings are consistent with prior work demonstrating a role for *grk-2* in *C. elegans* chemosensation [62–64]. Collectively, these findings show that VPA attraction requires at least one Gα protein, one RGS, and one GRK and is likely to be independent of arrestin, supporting a role for at least one GPCR serving as a molecular target for this drug.

## Discussion

Using *C. elegans* chemotaxis behavior as a screening tool, we showed that this simple animal is attracted to the anticonvulsant and mood-stabilizing drug, VPA. This whole-animal behavioral phenotype relies on chemosensation since smell- and taste-blind *tax-4;osm-9* double mutants are insensitive to VPA. We further show that VPA attraction requires an intact pair of AWC chemosensory neurons and that their contribution to behavior requires an intact *odr-1* and *gcy-28* rGC genes and the *tax-4* CNG-gated ion channel gene. The *tax-4* gene functions cell-autonomously within the AWC neurons. With *in vivo* chemosensory neuron-specific calcium imaging, we demonstrate that AWC activity is suppressed in the presence of VPA and exhibits a robust OFF response, reminiscent of the modulation of cytoplasmic calcium in AWC neurons stimulated by a number of other odorant ligands (e.g. Ref. [30,49,65–67]). All aspects of this VPA-evoked cellular response are eliminated by loss-of-function mutations in *tax-4* and in two rGC-encoding genes, *odr-1* and *gcy-28*. Collectively, these findings imply that VPA sensing depends on the activation of the AWC neurons and that VPA activates a cyclic nucleotide-mediated signaling cascade. Our genetic dissection studies implicate additional signaling partners in VPA attraction, including four genes encoding G protein L subunits, *odr-3*, *egl-30*, *gpa-2*, and *gpa-3*, two genes encoding regulators of G protein signaling (RGS) proteins, *rgs-2* and *rgs-3*, and one gene encoding a GPCR kinase (GRK), *grk-2*. All of these G protein related genes are expressed in the AWC neurons [46,47,54], marking them as candidate signaling partners accounting for VPA sensing by the AWC neurons. The picture emerging from these findings is that VPA attraction engages a CNG-mediated signaling cascade in the AWC neurons.

### Multiple chemosensory neurons detect VPA and contribute to attraction

The AWC neurons are not the only CSNs that contribute to VPA attraction or that generate CNG-sensitive calcium transients upon exposure to VPA. Loss of the ASH neurons increases VPA attraction (Fig. 2B), which is aligned with the tight association between ASH signaling and aversive responses [22]. Although the *odr-1* gene is not known to be expressed in the ASH neurons, loss of *odr-1* rGC function increased VPA- evoked calcium transients in the ASH neurons (Fig. 5). This effect might arise through conventional synaptic signaling or by negative regulation of ASH function mediated by cGMP transferred from *odr-1-*expressing neurons via gap junctions, as previously proposed for other ASH-dependent behaviors [68]. Three other pairs of chemosensory neurons showed *tax-4*-sensitive calcium responses: AWB, ADF, and ASK (Figure 3, Supplementary Figure 5). Similar to the AWC neurons, VPA-evoked calcium responses in AWB were absent in *tax-4*, *odr-1*, and *gcy-28* mutant animals (Figures 3, 5). Distinct from the AWC neurons, which showed robust VPA responses in every control animal, the AWB neurons responded to VPA in a subset of control animals. This variability might indicate that AWB is less sensitive to VPA than AWC, despite sharing a similar dependence upon *tax-4*, *odr-1*, and *gcy-28.* The engagement of multiple CSNs in VPA attraction and the unique genetic footprint of VPA-evoked calcium transients exhibited by each of the VPA-sensitive neurons (AWC, AWB, ASH) may indicate that each of these CSNs has a distinct VPA chemoreceptor(s), or that the sensitivity of the CNG signaling cascade differs among them, or a combination of these factors.

### The receptor guanylate cyclases required for VPA attraction contribute to sensing other chemicals

In addition to their role as enzymes that catalyze the synthesis of cGMP from GTP, rGCs also function as receptors for peptides and other chemical cues [22]. Prior studies in *C. elegans* implicate *odr-1* in chemotaxis responses to isoamyl alcohol, butanone, benzaldehyde, indole, and 2-heptanone [48,69,70] and *gcy-28* in butanone responses [50]. Consistent with the idea that these rGCs play a ligand-independent role in VPA chemotaxis, we found that *odr-1* is required for attraction not only to VPA, but also to isoamyl alcohol, furfural, 2-methyl-1-butanol, and valeric acid and that *gcy-28* is needed for attraction to VPA and 2-methyl-1-butanol. In addition to its expression in AWC, *odr-1* is expressed in ASI, ASJ, ASK, and AWB [24]. None of these CSN pairs responded to VPA as robustly as AWC in our calcium imaging experiments. Similarly, *gcy-28* transcripts are detected in most CSN pairs [46,47], but only a subset of CSNs are sensitive to VPA. Thus, neither *odr-1* nor *gcy-28* expression is insufficient by itself to confer sensitivity to VPA and both are required for attraction to multiple chemicals. Collectively, these findings imply that the contribution of *odr-1* and *gcy-28* to VPA sensing resides primarily in their enzymatic activity.

### VPA converges on a cGMP-gated ion channel and relies on cGMP synthesis

The AWC chemosensory neurons respond to chemical stimuli in a manner that is analogous to OFF responses in vertebrate photoreceptors [67], including the modulation of a cGMP-gated ion channel. The AWC neurons respond similarly to VPA (Fig. 2) as they do to other compounds and complex mixtures [30,65,67]. Here we show that ligand-induced decreases in intracellular Ca^2+^ and OFF responses in AWC require the *tax-4* CNG ion channel gene and the *odr-1* and *gcy-28* rGCs. These findings support a model in which TAX-4 ion channels are open in the absence of chemical stimuli and closed during chemical stimulation and that these changes in ion channel activity mirror intracellular cGMP concentrations. In this framework, VPA is predicted to reduce intracellular cGMP by inhibiting its synthesis or increasing its degradation (Fig. 6, top) and its removal increases intracellular cGMP (Fig. 6, bottom). We propose that the VPA-induced decrease in intracellular Ca^2+^ prompts a Ca^2+^-sensitive guanylate cyclase activating protein (GCAP) to activate rGCs, as found in vertebrate photoreceptors [reviewed by [53]]. Such a calcium-dependent feedback mechanism is a good candidate to account for an OFF response mediated by the hypothesized increase in intracellular cGMP and TAX-4-dependent ion channel activity.

**Figure 6:**
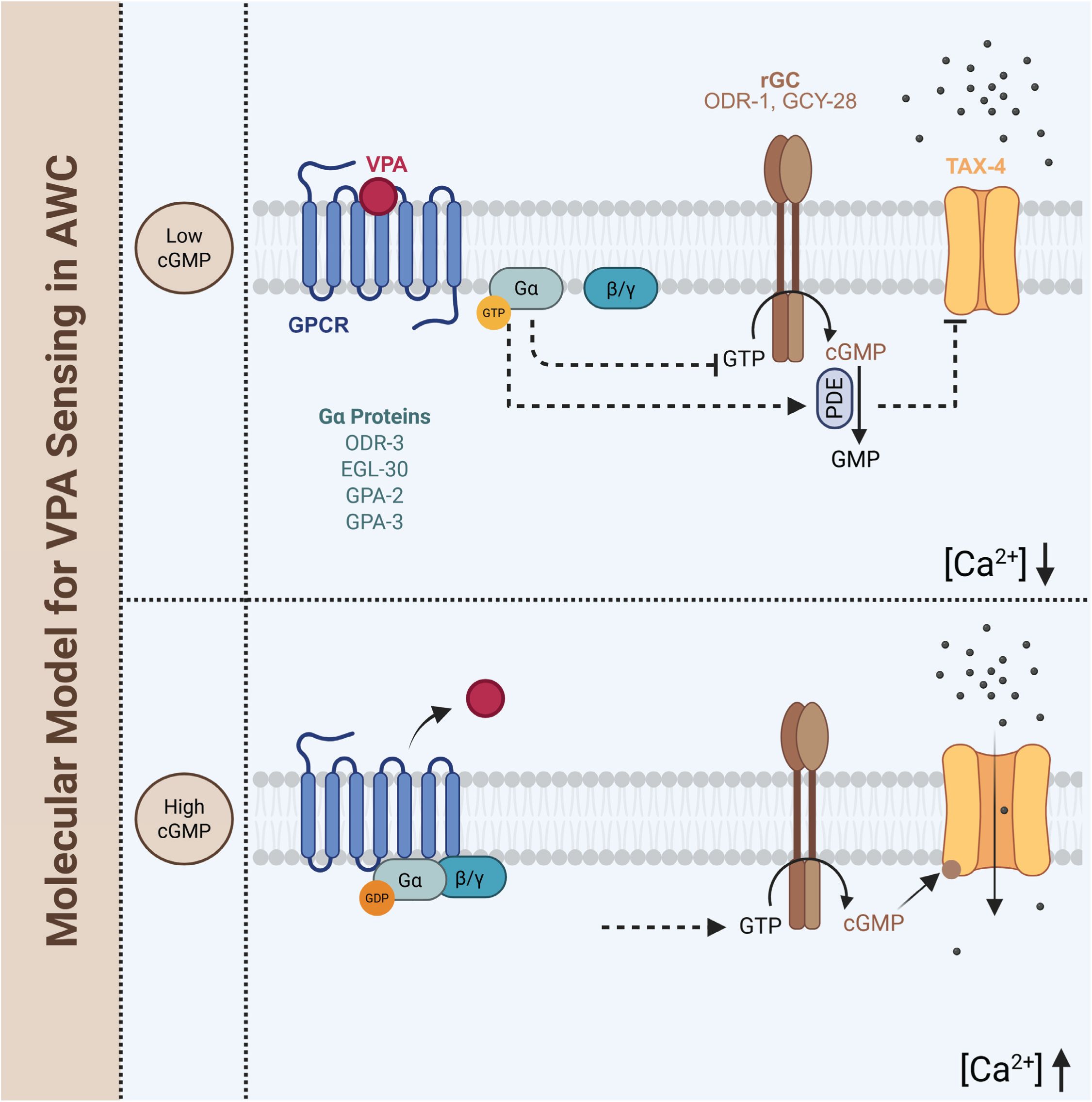
Molecular model for VPA sensing in AWC. **Low cGMP state** - VPA binds to and activates GPCR(s) present in the AWC neurons. This interaction results in the activation of its associated G protein, which leads to a decrease in cGMP levels inside the cell via its degradation by phosphodiesterases (PDEs) or the inhibition of its production. Consequently, lower cGMP levels result in the closing of *tax-4* ion channels, a reduction in Ca^2+^ ion influx, and hyperpolarization of the AWC neurons. **High cGMP state** - When VPA is removed or dissociates from its corresponding GPCR target(s), cGMP levels rise, *tax-4* ion channels open, cytoplasmic calcium levels increase, and the AWC neurons depolarize. Created in BioRender. Rogel-Hernandez, L. (2025) https://BioRender.com/15goo9i.

This proposed VPA signaling pathway is analogous to the canonical pathway underpinning light-dependent modulation of vertebrate photoreceptors and many of the proteins involved are conserved in *C. elegans* and in mammals, including humans. For instance, the *C. elegans* TAX-4 ion channel is orthologous to the human CNGA1 channel (51% identical) and the ODR-1 protein is orthologous to two human guanylate cyclases, GUCY2D (32% identical) and GUCY2F (34% identical). It remains to be determined whether the contribution of *odr-1* and *gcy-28* to VPA sensing is closer to the role played by rGCs in vertebrate phototransduction [53] or is instructive and modulated downstream of the GL proteins required for VPA attraction (Fig. 6, top). In either scenario, our finding that genes encoding GL proteins, RGS, and GRK proteins are essential for VPA attraction add evidence in favor of a role for a GPCR in VPA sensing. Although a VPA-sensitive GPCR has yet to be identified in any organism, members of this protein family are among the most pursued molecular targets for drug development and are often considered for repurposing due to their high druggability [71,72]. Identifying candidate GPCRs that bind VPA is complicated by the widespread finding that millimolar VPA (or that of its sodium salt analog, sodium valproate), is needed to modulate the activity of neurons in mammals [10–16] and other animals [7–9]. Our work helps to establish *C. elegans* as a model organism to study the mode of action of VPA in neurons and links VPA to cGMP-dependent signaling events. Whereas traditional approaches have so far failed to identify the low-affinity ligand-receptor interactions underpinning VPA’s modulation of brain activity, we have shown that a non-traditional approach grounded in the sensitivity of *C. elegans* chemotaxis and chemosensory transduction combined with genetic dissection is poised to overcome this barrier.

## Acknowledgments

We thank E. Fryer, C. Jaisinghani, and Z. Liao for assistance with coding, genetics, and laboratory management, respectively, and H. Lu for conducting pilot calcium imaging studies. Other members of the Goodman laboratory helped to improve data visualization and analysis and provided valuable feedback on writing. The following investigators and their laboratories contributed *C. elegans* strains: Chen (Umeå University), Y. Iino (The University of Tokyo), and P. Sengupta (Brandeis University), and several strains were provided by the CGC, which is funded by NIH Office of Research Infrastructure Programs (P40 OD010440). This work also benefited from Wormbase [60] in its design and execution. Work supported by grants from NIH to MBG (R35NS105092) and ADTS (U01NS132158), and a Stanford BioX Bowes Interdisciplinary Graduate Fellowship and NIH training grant (T32GM113854) to LERH.

---

**Supplementary Figure 1: Neural activity raster plots for all amphid CSN pairs in wild-type and *tax-4* mutant backgrounds.**

Neural activity (ΔF/F_0_) raster plots for all amphid CSN pairs (in alphabetical order) across control (A) and *tax-4* (B) calcium imaging lines. Within each raster plot, rows represent individual animals sorted by mean ΔF/F_0_ response during VPA presentation (5-15 seconds, region encapsulated by white dashed lines). The number of animals (*N*) contributing data for each CSN pair is displayed in the lower right corner. A shared colorbar legend can be found to the right that indicates the range of ΔF/F_0_.

**Supplementary Figure 2: The chemosensory neurons ASER, AWCL, and AWCR respond to VPA in a *tax-4*-dependent manner.**

**(A)** The neuronal activity of ASEL, ASER, AWCL, and AWCR neurons from control (navy blue) and *tax-4* (golden yellow) calcium imaging lines in response to VPA (6.12 × 10^-4^ M, 1:10^-4^ dilution) presentation and removal averaged across multiple trials (*N*). A shaded gray box indicates the 10-second VPA delivery period. Navy blue and golden yellow shaded areas represent the SEM for control and *tax-4*, respectively. **(B)** Neural activity (ΔF/F_0_) raster plots for ASEL, ASER, AWCL, and AWCR across control (Top) and *tax-4* (Bottom) calcium imaging lines. Within each raster plot, rows represent individual animals sorted by mean ΔF/F_0_ response during VPA presentation (5-15 seconds, region encapsulated by white dashed lines). The number of animals (*N*) contributing data for each CSN pair is displayed in the lower right corner. A shared colorbar legend can be found to the right that indicates the range of ΔF/F_0_.

**Supplementary Figure 3: The chemosensory neurons AWCL and AWCR respond to VPA in an *odr-1*- and *gcy-28*-dependent manner, whereas neither ASEL nor ASER do so.**

**(A)** Neural activity (ΔF/F_0_) raster plots for ASEL, ASER, AWCL, and AWCR across *odr-1* (Top) and *gcy-28* (Bottom) calcium imaging lines. Within each raster plot, rows represent individual animals sorted by mean ΔF/F_0_ response during VPA presentation (5-15 seconds, region encapsulated by white dashed lines). The number of animals (*N*) contributing data for each CSN pair is displayed in the lower right corner. A shared colorbar legend can be found to the right that indicates the range of ΔF/F_0_. **(B)** Quantification of the signed max ΔF/F_0_ value during VPA presentation for ASEL, ASER, AWCL, and AWCR across four calcium imaging lines: control, *tax-4*, *odr-1*, and *gcy-28*. Bars represent SEM and italicized numbers above tick marks in the x-axis indicate the number of trials (*N*) for each CSN pair. Statistically significant comparisons (*p* < 0.05) as a function of genotype in each neuron are annotated with an asterisk (Mann-Whitney U test).

**Supplementary Figure 4: Neural activity raster plots for all amphid CSN pairs in *odr-1* and *gcy-28* mutant backgrounds.**

Neural activity (ΔF/F_0_) raster plots for all amphid CSN pairs (in alphabetical order) across *odr-1* **(A)** and *gcy-28* **(B)** calcium imaging lines. Within each raster plot, rows represent individual animals sorted by mean ΔF/F_0_ response during VPA presentation (5-15 seconds, region encapsulated by white dashed lines). The number of animals (*N*) contributing data for each CSN pair is displayed in the lower right corner. A shared colorbar legend can be found to the right that indicates the range of ΔF/F_0_.

**Supplementary Figure 5: Neuronal activity of eight CSN pairs in *odr-1* and *gcy-28* mutant backgrounds that display small or no responses to VPA in wild-type.**

**(A)** The neuronal activity of eight CSNs (ADF, ADL, ASE, ASG, ASI, ASJ, ASK, and AWA) from *odr-1* (dark green) and *gcy-28* (burnt orange) calcium imaging lines in response to VPA (6.12 × 10^-4^ M, 1:10^-4^ dilution) presentation and removal averaged across multiple trials (*N*). A shaded gray box indicates the 10-second VPA delivery period. Dark green and burnt orange shaded areas represent the SEM for *odr-1* and *gcy-28*, respectively. **(B)** Quantification of the signed max ΔF/F_0_ value during VPA presentation for ADF, ADL, ASE, ASG, ASI, ASJ, ASK, and AWA across four calcium imaging lines: control, *tax-4*, *odr-1*, and *gcy-28*. Bars represent SEM and italicized numbers above tick marks in the x-axis indicate the number of trials (*N*) for each CSN pair. Statistically significant comparisons (*p* < 0.05) as a function of genotype in each neuron are annotated with an asterisk (Mann-Whitney U test).

## Notes

### Competing Interest Statement

The authors have declared no competing interest.

### Summary of Updates

Text and citations revised to be more concise; added a new discussion of prior studies of gcy-28 mutants in chemosensation.

