## Supplementary Table 1 for "The anticonvulsant and mood-stabilizing drug valproic acid attracts *C. elegans* and activates chemosensory neurons via a cGMP signaling pathway"

| *C. elegans* strain | Genotype | Source |
|---|---|---|
| CHS2307 | *gcy-12(yum88)* II | Chen Lab |
| CHS502 | *gcy-28(yum32)* I | Chen Lab |
| CX10 | *osm-9(ky10)* IV | CGC |
| CX2065 | *odr-1(n1936)* X | CGC |
| CX4 | *odr-7(ky4)* X | CGC |
| CX5757 | *kyIs140* I*; nsy-4(ky616)* IV | GCG |
| CX7102 | *qaIs2241 [gcy-36::egl-1 + gcy-35::GFP + lin-15(+)]* | CGC |
| DA1084 | *egl-30(ad806)* I | CGC |
| DG1856 | *goa-1(sa734)* I | CGC |
| FG7 | *grk-2(gk268)* III | CGC |
| GN1077 | *tax-4(p678)* III*;osm-9(ky10)* IV | In-house |
| GN1103 | *tax-4(p678)* III*; aezaIs008 [ift-20p::GCaMP6s::3x::NLS]* ?*; hpIs728 [gpc-1p::mCherry]* X | This study |
| GN1126 | *odr-1(n1936)* X*; aezaIs008 [ift-20p::GCaMP6s::3x::NLS]* ?*; hpIs728 [gpc-1p::mCherry]* X | This study |
| GN1136 | *gcy-28(yum32)* I*; aezaIs008 [ift-20p::GCaMP6s::3x::NLS]* ?*; hpIs728 [gpc-1p::mCherry]* X | This study |
| GN576 | *tax-4(p678)* III*; pgIs17[ceh-36p::tax-4(+) + unc-122p::DsRed]* | In-house |
| GN579 | *tax-4(p678)* III*; pgIs18[srbc-65p::tax-4(+) + unc-122p::DsRed]* | In-house |
| GN580 | *pgIs19(jxEx100(pQZ37(trx-1::ICE);ofm-1::GFP))* | In-house |
| JN1194 | *gcy-14(pe1102)* V | CGC |
| JN1713 | *peIs1713 [sra-6p::mCasp-1 + unc-122p::mCherry]* | Iino Lab |
| JN1715 | *peIs1715[str-1p::mCasp1 + unc-122p::GFP]* | Iino Lab |
| JN2113 | *peIs2113 [gcy-21p::mCaspase + tax-4p::NLS::YC2.60 + lin-44p::GFP]* | Iino Lab |
| LX160 | *rgs-2(vs17)* X | CGC |
| MT4810 | *odr-3(n2046)* V | CGC |
| N2 | Wild-type | CGC |
| NL1137 | *gpa-5(pk376)* X | CGC |
| NL2330 | *gpa-13(pk1270)* V | CGC |
| NL334 | *gpa-2(pk16)* V | CGC |
| NL335 | *gpa-3(pk35)* V | CGC |
| NL348 | *gpa-2(pk16);gpa-3(pk35)* V | CGC |
| NL795 | *gpa-7(pk610)* IV | CGC |
| OH13098 | *che-1(ot75)* I | CGC |
| OH13813 | *oig-8(ot818)* II | CGC |
| PR678 | *tax-4(p678)* III | CGC |
| PS6025 | *qrIs2[sra-9::mCasp1]* | CGC |
| PY7502 | *oyIs85 [ceh-36p::TU#813 + ceh-36p::TU#814 + srtx-1p::GFP + unc-122p::DsRed]* | Sengupta Lab |
| PY7505 | *oyIs84 [gpa-4p::TU#813 + gcy-27p::TU#814 + gcy-27p::GFP + unc-122p::DsRed]* | Sengupta Lab |
| RB1194 | *grk-1(ok1239)* X | CGC |
| RB1780 | *rgs-3(ok2288)* II | CGC |
| RB1816 | *gpa-16(ok2349)* I | CGC |
| RB1935 | *gcy-20(ok2538)* V | CGC |
| RB660 | *arr-1(ok401)* X | CGC |
| SP1735 | *dyf-7(m537)* X | CGC |
| VC390 | *nsy-1(ok593)* II | CGC |
| ZM10104 | *aezaIs008 [ift-20p::GCaMP6s::3x::NLS]* ?*; hpIs728 [gpc-1p::mCherry]* X | Samuel Lab |
