## Supplementary Table 2 for "The anticonvulsant and mood-stabilizing drug valproic acid attracts *C. elegans* and activates chemosensory neurons via a cGMP signaling pathway"

| Figure | Compound | Concentration (M) | Genetic Background | Mean Worm Location (mm) | Standard Deviation (± mm) |
|---|---|---|---|---|---|
| 1B | Water | Pure | WT | 1.74 | 17.08 |
| 1B | VPA | 0.0204 | WT | 1.17 | 16.98 |
| 1B | VPA | 0.0612 | WT | 1.80 | 16.30 |
| 1B | VPA | 0.204 | WT | 4.10 | 16.99 |
| 1B | VPA | 0.612 | WT | 5.16 | 16.96 |
| 1B | VPA | 6.12 | WT | 6.82 | 15.29 |
| 1D | VPA | 6.12 | WT | 7.83 | 14.65 |
| 1D | VPA | 6.12 | *dyf-7* | 2.04 | 13.29 |
| 1D | VPA | 6.12 | *oig-8* | 4.79 | 15.07 |
| 1E | VPA | 6.12 | WT | 7.83 | 14.65 |
| 1E | VPA | 6.12 | *tax-4;osm-9* | 1.69 | 16.43 |
| 1E | VPA | 6.12 | *osm-9* | 11.28 | 13.50 |
| 1E | VPA | 6.12 | *tax-4* | -2.22 | 13.21 |
| 2B | VPA | 6.12 | WT | 7.83 | 14.65 |
| 2B | VPA | 6.12 | *tax-4* | -2.22 | 13.21 |
| 2B | VPA | 6.12 | ASH- | 10.16 | 14.90 |
| 2B | VPA | 6.12 | *odr-7* | 8.88 | 15.45 |
| 2B | VPA | 6.12 | ASK- | 7.26 | 15.52 |
| 2B | VPA | 6.12 | AWB- | 7.09 | 15.12 |
| 2B | VPA | 6.12 | *che-1* | 7.09 | 14.84 |
| 2B | VPA | 6.12 | ASI- | 6.76 | 13.46 |
| 2B | VPA | 6.12 | ASJ- | 6.29 | 14.48 |
| 2B | VPA | 6.12 | ASG- | 6.02 | 15.10 |
| 2B | VPA | 6.12 | A/P/U- | 5.14 | 15.90 |
| 2B | VPA | 6.12 | AWC- | 1.08 | 14.96 |
| 2C | VPA | 6.12 | WT | 7.83 | 14.65 |
| 2C | VPA | 6.12 | *tax-4* | -2.22 | 13.21 |
| 2C | VPA | 6.12 | *tax-4* rescue in ASI | -0.82 | 13.11 |
| 2C | VPA | 6.12 | *tax-4* rescue in AWC | 7.82 | 14.99 |
| 2D | VPA | 6.12 | WT | 7.83 | 14.65 |
| 2D | VPA | 6.12 | AWC- | 1.08 | 14.96 |
| 2D | VPA | 6.12 | *nsy-1* | 3.27 | 15.97 |
| 2D | VPA | 6.12 | *nsy-4* | 2.97 | 15.00 |
| 4B | VPA | 6.12 | WT | 7.83 | 14.65 |
| 4B | VPA | 6.12 | *gcy-12* | 9.40 | 15.00 |
| 4B | VPA | 6.12 | *gcy-14* | 8.92 | 15.21 |
| 4B | VPA | 6.12 | *gcy-20* | 6.20 | 15.92 |
| 4B | VPA | 6.12 | *odr-1* | 1.45 | 16.12 |
| 4B | VPA | 6.12 | *gcy-28* | -0.31 | 16.15 |
| 4C | 2M1B | 0.02 | WT | 8.38 | 15.01 |
| 4C | 2M1B | 0.02 | *tax-4* | 0.29 | 16.14 |

| Figure | Compound | Concentration (M) | Genetic Background | Mean Worm Location (mm) | Standard Deviation (± mm) |
|---|---|---|---|---|---|
| 4C | 2M1B | 0.02 | *odr-1* | 2.50 | 16.92 |
| 4C | 2M1B | 0.02 | *gcy-28* | 4.32 | 16.28 |
| 4C | FUR | 0.02 | WT | 5.46 | 16.47 |
| 4C | FUR | 0.02 | *tax-4* | 0.75 | 15.47 |
| 4C | FUR | 0.02 | *odr-1* | 1.62 | 17.45 |
| 4C | FUR | 0.02 | *gcy-28* | 5.82 | 16.81 |
| 4C | IAA | 0.0932 | WT | 7.99 | 13.93 |
| 4C | IAA | 0.0932 | *tax-4* | 0.35 | 14.93 |
| 4C | IAA | 0.0932 | *odr-1* | -0.14 | 17.00 |
| 4C | IAA | 0.0932 | *gcy-28* | 8.80 | 14.85 |
| 4C | VA | 9.1 | WT | 8.04 | 14.34 |
| 4C | VA | 9.1 | *tax-4* | -0.61 | 15.32 |
| 4C | VA | 9.1 | *odr-1* | -1.70 | 15.56 |
| 4C | VA | 9.1 | *gcy-28* | 8.63 | 13.68 |
| 4C | VPA | 6.12 | WT | 7.23 | 14.46 |
| 4C | VPA | 6.12 | *tax-4* | -0.78 | 15.03 |
| 4C | VPA | 6.12 | *odr-1* | 1.53 | 15.52 |
| 4C | VPA | 6.12 | *gcy-28* | 2.21 | 14.79 |
| 5C | VPA | 6.12 | WT | 7.83 | 14.65 |
| 5C | VPA | 6.12 | *gpa-2* | 11.24 | 12.74 |
| 5C | VPA | 6.12 | *gpa-7* | 7.33 | 14.24 |
| 5C | VPA | 6.12 | *gpa-16* | 6.86 | 15.42 |
| 5C | VPA | 6.12 | *gpa-5* | 6.10 | 14.13 |
| 5C | VPA | 6.12 | *gpa-3* | 5.51 | 14.98 |
| 5C | VPA | 6.12 | *goa-1* | 5.17 | 14.58 |
| 5C | VPA | 6.12 | *gpa-13* | 4.69 | 15.16 |
| 5C | VPA | 6.12 | *odr-3* | 3.22 | 15.97 |
| 5C | VPA | 6.12 | *egl-30* | 2.69 | 14.63 |
| 5C | VPA | 6.12 | *gpa-2;gpa-3* | 2.30 | 16.42 |
| 5D | VPA | 6.12 | WT | 7.83 | 14.65 |
| 5D | VPA | 6.12 | *rgs-2* | 4.78 | 15.63 |
| 5D | VPA | 6.12 | *rgs-3* | 2.72 | 14.11 |
| 5E | VPA | 6.12 | WT | 7.83 | 14.65 |
| 5E | VPA | 6.12 | *arr-1* | 7.41 | 13.48 |
| 5E | VPA | 6.12 | *grk-1* | 8.51 | 14.12 |
| 5E | VPA | 6.12 | *grk-2* | 4.27 | 15.27 |
