## Supplementary Table 3 for "The anticonvulsant and mood-stabilizing drug valproic acid attracts *C. elegans* and activates chemosensory neurons via a cGMP signaling pathway"

| Figure | Control | Test | Compound | Concentration (M) | <ΔLocation> (mm) | Lower 95% CI (mm) | Upper 95% CI (mm) |
|---|---|---|---|---|---|---|---|
| 1B | Water | VPA | VPA | 0.0204 | -0.57 | -1.32 | 0.20 |
| 1B | Water | VPA | VPA | 0.0612 | 0.06 | -0.68 | 0.82 |
| 1B | Water | VPA | VPA | 0.204 | 2.35 | 1.60 | 3.13 |
| 1B | Water | VPA | VPA | 0.612 | 3.42 | 2.58 | 4.20 |
| 1B | Water | VPA | VPA | 6.12 | 5.08 | 4.29 | 5.85 |
| 1D | WT | *dyf-7* | VPA | 6.12 | -5.79 | -6.47 | -5.09 |
| 1D | WT | *oig-8* | VPA | 6.12 | -3.04 | -3.72 | -2.38 |
| 1E | WT | *tax-4;osm-9* | VPA | 6.12 | -6.15 | -6.90 | -5.43 |
| 1E | WT | *osm-9* | VPA | 6.12 | 3.44 | 2.75 | 4.13 |
| 1E | WT | *tax-4* | VPA | 6.12 | -10.05 | -10.78 | -9.36 |
| 2B | WT | *tax-4* | VPA | 6.12 | -10.05 | -10.78 | -9.36 |
| 2B | WT | ASH- | VPA | 6.12 | 2.33 | 1.58 | 3.14 |
| 2B | WT | *odr-7* | VPA | 6.12 | 1.05 | 0.27 | 1.87 |
| 2B | WT | ASK- | VPA | 6.12 | -0.57 | -1.37 | 0.17 |
| 2B | WT | AWB- | VPA | 6.12 | -0.74 | -1.54 | 0.06 |
| 2B | WT | *che-1* | VPA | 6.12 | -0.74 | -1.56 | 0.11 |
| 2B | WT | ASI- | VPA | 6.12 | -1.07 | -1.90 | -0.32 |
| 2B | WT | ASJ- | VPA | 6.12 | -1.54 | -2.26 | -0.84 |
| 2B | WT | ASG- | VPA | 6.12 | -1.82 | -2.54 | -1.13 |
| 2B | WT | A/P/U- | VPA | 6.12 | -2.70 | -3.48 | -1.93 |
| 2B | WT | AWC- | VPA | 6.12 | -6.76 | -7.37 | -6.16 |
| 2C | WT | *tax-4* | VPA | 6.12 | -10.05 | -10.78 | -9.36 |
| 2C | WT | *tax-4* rescue in ASI | VPA | 6.12 | -8.65 | -9.24 | -8.05 |
| 2C | WT | *tax-4* rescue in AWC | VPA | 6.12 | -0.02 | -0.85 | 0.79 |
| 2D | WT | AWC- | VPA | 6.12 | -6.76 | -7.37 | -6.16 |
| 2D | WT | *nsy-1* | VPA | 6.12 | -4.56 | -5.39 | -3.72 |
| 2D | WT | *nsy-4* | VPA | 6.12 | -4.86 | -5.64 | -4.09 |
| 4B | WT | *gcy-12* | VPA | 6.12 | 1.56 | 0.81 | 2.28 |
| 4B | WT | *gcy-14* | VPA | 6.12 | 1.09 | 0.37 | 1.81 |
| 4B | WT | *gcy-20* | VPA | 6.12 | -1.64 | -2.38 | -0.96 |
| 4B | WT | *odr-1* | VPA | 6.12 | -6.38 | -7.09 | -5.69 |
| 4B | WT | *gcy-28* | VPA | 6.12 | -8.15 | -8.80 | -7.47 |
| 4C | WT | *tax-4* | 2M1B | 0.02 | -8.09 | -8.83 | -7.35 |
| 4C | WT | *odr-1* | 2M1B | 0.02 | -5.88 | -6.57 | -5.12 |
| 4C | WT | *gcy-28* | 2M1B | 0.02 | -4.06 | -4.88 | -3.28 |
| 4C | WT | *tax-4* | FUR | 0.02 | -4.70 | -5.56 | -3.84 |
| 4C | WT | *odr-1* | FUR | 0.02 | -3.83 | -4.75 | -2.88 |
| 4C | WT | *gcy-28* | FUR | 0.02 | 0.37 | -0.54 | 1.25 |
| 4C | WT | *tax-4* | IAA | 0.0932 | -7.64 | -8.26 | -6.97 |
| 4C | WT | *odr-1* | IAA | 0.0932 | -8.13 | -8.85 | -7.42 |
| 4C | WT | *gcy-28* | IAA | 0.0932 | 0.81 | 0.11 | 1.52 |

| Figure | Control | Test | Compound | Concentration (M) | <ΔLocation> (mm) | Lower 95% CI (mm) | Upper 95% CI (mm) |
|---|---|---|---|---|---|---|---|
| 4C | WT | *tax-4* | VA | 9.1 | -8.64 | -9.43 | -7.85 |
| 4C | WT | *odr-1* | VA | 9.1 | -9.74 | -10.43 | -9.05 |
| 4C | WT | *gcy-28* | VA | 9.1 | 0.60 | -0.11 | 1.31 |
| 4C | WT | *tax-4* | VPA | 6.12 | -8.01 | -8.59 | -7.43 |
| 4C | WT | *odr-1* | VPA | 6.12 | -5.71 | -6.39 | -5.03 |
| 4C | WT | *gcy-28* | VPA | 6.12 | -5.03 | -5.61 | -4.42 |
| 5C | WT | *gpa-2* | VPA | 6.12 | 3.41 | 2.63 | 4.11 |
| 5C | WT | *gpa-7* | VPA | 6.12 | -0.50 | -1.23 | 0.24 |
| 5C | WT | *gpa-16* | VPA | 6.12 | -0.98 | -1.82 | -0.09 |
| 5C | WT | *gpa-5* | VPA | 6.12 | -1.73 | -2.48 | -1.02 |
| 5C | WT | *gpa-3* | VPA | 6.12 | -2.32 | -2.95 | -1.68 |
| 5C | WT | *goa-1* | VPA | 6.12 | -2.66 | -3.33 | -2.03 |
| 5C | WT | *gpa-13* | VPA | 6.12 | -3.14 | -3.85 | -2.41 |
| 5C | WT | *odr-3* | VPA | 6.12 | -4.61 | -5.29 | -3.99 |
| 5C | WT | *egl-30* | VPA | 6.12 | -5.14 | -5.92 | -4.35 |
| 5C | WT | *gpa-2;gpa-3* | VPA | 6.12 | -5.53 | -6.18 | -4.88 |
| 5D | WT | *rgs-2* | VPA | 6.12 | -3.05 | -3.75 | -2.37 |
| 5D | WT | *rgs-3* | VPA | 6.12 | -5.12 | -5.86 | -4.40 |
| 5E | WT | *arr-1* | VPA | 6.12 | -0.43 | -1.13 | 0.29 |
| 5E | WT | *grk-1* | VPA | 6.12 | 0.67 | -0.07 | 1.38 |
| 5E | WT | *grk-2* | VPA | 6.12 | -3.56 | -4.33 | -2.81 |
