## Supplementary Table 4 for "The anticonvulsant and mood-stabilizing drug valproic acid attracts *C. elegans* and activates chemosensory neurons via a cGMP signaling pathway"

| CSN | Calcium Imaging Line | Mean Signed Max ΔF/F0 | SEM |
|---|---|---|---|
| ADF | Control | -0.00704341 | 0.007917806 |
| ADL | Control | -0.006509483 | 0.005750596 |
| ASE | Control | 0.000524106 | 0.029326457 |
| ASG | Control | -0.014027376 | 0.004882591 |
| ASH | Control | 0.028435251 | 0.00962705 |
| ASI | Control | -0.005450313 | 0.007480756 |
| ASJ | Control | -0.023283077 | 0.007009967 |
| ASK | Control | -0.040173013 | 0.004862621 |
| AWA | Control | -0.021312892 | 0.0031788 |
| AWB | Control | 0.04290873 | 0.037912511 |
| AWC | Control | -0.164593586 | 0.026988708 |
| ASEL | Control | -0.022166938 | 0.00627127 |
| ASER | Control | -0.000385185 | 0.050220108 |
| AWCL | Control | -0.190984918 | 0.030394384 |
| AWCR | Control | -0.128807363 | 0.049547753 |
| ADF | *gcy-28* | -0.044327869 | 0.007801474 |
| ADL | *gcy-28* | -0.02935617 | 0.009966114 |
| ASE | *gcy-28* | -0.015730737 | 0.015635369 |
| ASG | *gcy-28* | -0.01291874 | 0.01039367 |
| ASH | *gcy-28* | 0.043342865 | 0.024238467 |
| ASI | *gcy-28* | -0.013193425 | 0.005700443 |
| ASJ | *gcy-28* | -0.024353536 | 0.009329265 |
| ASK | *gcy-28* | -0.019952283 | 0.009366861 |
| AWA | *gcy-28* | -0.026310397 | 0.003426799 |
| AWB | *gcy-28* | 0.025309854 | 0.023422558 |
| AWC | *gcy-28* | -0.002631101 | 0.016961313 |
| ASEL | *gcy-28* | 0.028345342 | 0.04166015 |
| ASER | *gcy-28* | -0.018533688 | 0.021017874 |
| AWCL | *gcy-28* | -0.001323194 | 0.02810517 |
| AWCR | *gcy-28* | -0.00460944 | 0.015725188 |
| ADF | *odr-1* | -0.016176787 | 0.006497253 |
| ADL | *odr-1* | -0.0171373 | 0.007847278 |
| ASE | *odr-1* | -0.03796478 | 0.009979313 |
| ASG | *odr-1* | -0.018603376 | 0.006896922 |
| ASH | *odr-1* | 0.172218726 | 0.059621424 |
| ASI | *odr-1* | -0.02644007 | 0.004594344 |
| ASJ | *odr-1* | -0.030130545 | 0.016449023 |
| ASK | *odr-1* | -0.082641848 | 0.027033627 |
| AWA | *odr-1* | 0.018339995 | 0.030128292 |
| AWB | *odr-1* | 0.001911984 | 0.010223065 |
| AWC | *odr-1* | 0.012431951 | 0.012213573 |

| CSN | Calcium Imaging Line | Mean Signed Max ΔF/F0 | SEM |
|---|---|---|---|
| ASEL | *odr-1* | -0.023017481 | 0.013745504 |
| ASER | *odr-1* | -0.01873454 | 0.037375389 |
| AWCL | *odr-1* | 0.035461186 | 0.014098336 |
| AWCR | *odr-1* | 0.002917723 | 0.015796875 |
| ADF | *tax-4* | -0.022125734 | 0.003897231 |
| ADL | *tax-4* | -0.013501967 | 0.003483753 |
| ASE | *tax-4* | -0.01103871 | 0.006418173 |
| ASG | *tax-4* | -0.008532755 | 0.004734906 |
| ASH | *tax-4* | 0.038377476 | 0.018201104 |
| ASI | *tax-4* | -0.016529488 | 0.00196109 |
| ASJ | *tax-4* | -0.018593985 | 0.001680996 |
| ASK | *tax-4* | -0.017429263 | 0.003205883 |
| AWA | *tax-4* | -0.015133253 | 0.006623627 |
| AWB | *tax-4* | -0.009799881 | 0.014471021 |
| AWC | *tax-4* | 0.002780434 | 0.00642781 |
| ASEL | *tax-4* | -0.007194513 | 0.008616009 |
| ASER | *tax-4* | -0.018493117 | 0.004051152 |
| AWCL | *tax-4* | 0.002128726 | 0.009599246 |
| AWCR | *tax-4* | 0.006124261 | 0.010059143 |
