## Supplementary Table 5 for "The anticonvulsant and mood-stabilizing drug valproic acid attracts *C. elegans* and activates chemosensory neurons via a cGMP signaling pathway"

| CSN | Group 1 | Group 2 | Group 1 ($N$) | Group 2 ($N$) | Group 1 Median | Group 2 Median | U-statistic | $p$ -value |
|---|---|---|---|---|---|---|---|---|
| ADF | *tax-4* | Control | 10 | 16 | -0.024139 | -0.014809 | 40 | 0.037358 |
| ADL | *tax-4* | Control | 11 | 19 | -0.015702 | -0.011151 | 72 | 0.168461 |
| ASE | *tax-4* | Control | 12 | 17 | -0.019333 | -0.028057 | 135 | 0.150115 |
| ASEL | *tax-4* | Control | 12 | 15 | -0.017470 | -0.029447 | 116 | 0.213399 |
| ASER | *tax-4* | Control | 10 | 12 | -0.022324 | -0.071993 | 74 | 0.373378 |
| ASG | *tax-4* | Control | 12 | 13 | -0.011950 | -0.018721 | 92 | 0.462764 |
| ASH | *tax-4* | Control | 14 | 19 | 0.007802 | 0.029713 | 135 | 0.956426 |
| ASI | *tax-4* | Control | 11 | 18 | -0.016431 | -0.014162 | 83 | 0.486007 |
| ASJ | *tax-4* | Control | 13 | 19 | -0.017329 | -0.029905 | 158 | 0.192043 |
| ASK | *tax-4* | Control | 11 | 19 | -0.019441 | -0.037166 | 168 | 0.006702 |
| AWA | *tax-4* | Control | 10 | 11 | -0.017351 | -0.014791 | 61 | 0.698535 |
| AWB | *tax-4* | Control | 13 | 13 | -0.017227 | -0.015132 | 76 | 0.681618 |
| AWC | *tax-4* | Control | 20 | 18 | -0.010607 | -0.178398 | 334 | 0.000007 |
| AWCL | *tax-4* | Control | 19 | 11 | -0.014615 | -0.183681 | 209 | 0.000008 |
| AWCR | *tax-4* | Control | 16 | 14 | -0.009656 | -0.146942 | 195 | 0.000605 |
| ADF | *odr-1* | Control | 7 | 16 | -0.016874 | -0.014809 | 47 | 0.570082 |
| ADL | *odr-1* | Control | 7 | 19 | -0.016887 | -0.011151 | 45 | 0.224765 |
| ASE | *odr-1* | Control | 6 | 17 | -0.029080 | -0.028057 | 44 | 0.649044 |
| ASEL | *odr-1* | Control | 6 | 15 | -0.030764 | -0.029447 | 43 | 0.907038 |
| ASER | *odr-1* | Control | 5 | 12 | -0.012480 | -0.071993 | 33 | 0.792147 |
| ASG | *odr-1* | Control | 7 | 13 | -0.021782 | -0.018721 | 34 | 0.383387 |
| ASH | *odr-1* | Control | 7 | 19 | 0.218618 | 0.029713 | 99 | 0.064337 |
| ASI | *odr-1* | Control | 7 | 18 | -0.024261 | -0.014162 | 21 | 0.012015 |
| ASJ | *odr-1* | Control | 6 | 19 | -0.032584 | -0.029905 | 52 | 0.774627 |
| ASK | *odr-1* | Control | 7 | 19 | -0.081395 | -0.037166 | 34 | 0.064337 |
| AWA | *odr-1* | Control | 7 | 11 | -0.014684 | -0.014791 | 50 | 0.319137 |
| AWB | *odr-1* | Control | 7 | 13 | 0.006618 | -0.015132 | 47 | 0.936839 |
| AWC | *odr-1* | Control | 7 | 18 | 0.025957 | -0.178398 | 117 | 0.001204 |
| AWCL | *odr-1* | Control | 6 | 11 | 0.034012 | -0.183681 | 66 | 0.001089 |
| AWCR | *odr-1* | Control | 6 | 14 | 0.005697 | -0.146942 | 73 | 0.011883 |
| ADF | *gcy-28* | Control | 3 | 16 | -0.041596 | -0.014809 | 1 | 0.011884 |
| ADL | *gcy-28* | Control | 5 | 19 | -0.021340 | -0.011151 | 15 | 0.022929 |
| ASE | *gcy-28* | Control | 4 | 17 | -0.029743 | -0.028057 | 36 | 0.893131 |
| ASEL | *gcy-28* | Control | 3 | 15 | 0.030524 | -0.029447 | 33 | 0.236137 |
| ASER | *gcy-28* | Control | 4 | 12 | -0.025115 | -0.071993 | 29 | 0.585269 |
| ASG | *gcy-28* | Control | 4 | 13 | -0.021106 | -0.018721 | 25 | 0.954853 |
| ASH | *gcy-28* | Control | 5 | 19 | 0.042321 | 0.029713 | 55 | 0.618785 |
| ASI | *gcy-28* | Control | 5 | 18 | -0.012840 | -0.014162 | 45 | 0.970271 |
| ASJ | *gcy-28* | Control | 5 | 19 | -0.032746 | -0.029905 | 48 | 1.000000 |
| ASK | *gcy-28* | Control | 5 | 19 | -0.027122 | -0.037166 | 68 | 0.155132 |
| AWA | *gcy-28* | Control | 3 | 11 | -0.028353 | -0.014791 | 11 | 0.436275 |
| AWB | *gcy-28* | Control | 4 | 13 | 0.016744 | -0.015132 | 30 | 0.691886 |
| AWC | *gcy-28* | Control | 5 | 18 | -0.007209 | -0.178398 | 82 | 0.006517 |

| CSN | Group 1 | Group 2 | Group 1 ($N$) | Group 2 ($N$) | Group 1 Median | Group 2 Median | U-statistic | $p$ -value |
|---|---|---|---|---|---|---|---|---|
| AWCL | *gcy-28* | Control | 3 | 11 | 0.015699 | -0.183681 | 32 | 0.019517 |
| AWCR | *gcy-28* | Control | 5 | 14 | -0.012264 | -0.146942 | 60 | 0.023313 |
